# Multiplatform molecular profiling and functional genomic screens identify prognostic signatures and mechanisms underlying MEK inhibitor response in somatic *NF1* mutant glioblastoma

**DOI:** 10.1101/2024.07.01.601334

**Authors:** Sixuan Pan, Kanish Mirchia, Emily Payne, S. John Liu, Nadeem Al-Adli, Zain Peeran, Poojan Shukla, Jacob S. Young, Rohit Gupta, Jasper Wu, Joanna Pak, Kyounghee Seo, Tomoko Ozawa, Brian Na, Alyssa T. Reddy, Steve E. Braunstein, Joanna J. Phillips, Susan Chang, David A. Solomon, Arie Perry, David R. Raleigh, Mitchel S. Berger, Adam R. Abate, Harish N. Vasudevan

## Abstract

*NF1* is recurrently mutated in glioblastoma yet the molecular landscape and efficacy of targeted therapies remain unclear. Here, we combine bulk and single cell genomics of human somatic *NF1* mutant, IDH-wildtype glioblastomas with functional genomic analysis of cell lines and mouse intracranial tumor models to identify molecular subgroups within *NF1* mutant glioblastomas and mechanisms underlying MEK inhibitor response. Targeted DNA sequencing showed homozygous deletion of the cell cycle regulator *CDKN2A/B* is a poor prognostic marker in somatic *NF1* mutant, but not *NF1* wildtype, tumors. DNA methylation array profiling revealed three epigenetic groups highlighted by distinct clinical features, co-mutation patterns, and reference methylation classifier identities. Genome-wide CRISPRi screens in glioblastoma cells revealed cell cycle regulators are conserved mediators of cell growth while response to the MEK inhibitor selumetinib converges on Ras/RAF/MEK pathway activation. Repression of the RAF regulator *SHOC2* sensitizes glioblastomas to selumetinib *in vitro* and *in vivo* in mouse intracranial glioblastoma models. Single cell RNA-sequencing of mouse intracranial glioblastomas treated with the MEK inhibitor selumetinib reveals distinct responses between mesenchymal-like (MES-like) and non MES-like subpopulations suggesting non-MES like cells are intrinsically resistant to MEK inhibition. Finally, single nuclear RNA-sequencing (snRNA-seq) of human *NF1* mutant, *CDKN2A/B* deleted glioblastomas reveals MES-like tumor cells are associated with selumetinib sensitivity signatures while non MES-like cells exhibit increased cell cycle progression and lack selumetinib sensitivity, further supporting the notion that MEK inhibition specifically targets MES-like tumor cell subpopulations. Taken together, our data underscores the heterogeneity between and within somatic *NF1* mutant glioblastomas and delineates mechanisms of MEK inhibitor response across distinct tumor subpopulations, guiding the development of future therapeutic strategies that may synergize with MEK inhibition for *NF1* mutant tumors.

The tumor suppressor *NF1* is mutated in 15% of glioblastomas,^1–3^ the most common malignant brain tumor with poor outcomes and few effective treatments.^4^ NF1 is a GTPase activating protein (GAP) that negatively regulates Ras, and thus, *NF1* loss leads to induction of Ras/RAF/MEK/ERK signaling, driving tumorigenesis and comprising a targetable molecular cascade.^5,6^ Genomic analysis of glioblastoma demonstrates *NF1* mutation is associated with a mesenchymal-like (MES-like) transcriptomic tumor cell subpopulation and altered tumor microenvironment.^7,8^ More broadly, DNA methylation analysis reveals multiple epigenetic subgroups with overlapping relationships to transcriptomic subtype and DNA alterations, underscoring the complex relationship between genetic drivers and molecular signatures.^9^ While the updated 2021 Central Nervous System WHO tumor classification incorporates an ever increasing amount of molecular criteria for diffuse astrocytic tumors,^10^ the existence and clinical significance of molecular subgroups within somatic *NF1* mutant, IDH-wildtype glioblastomas based on genetic co-mutations, epigenetic profile, or transcriptomic signatures remain unclear.

Preclinical data support the utility of MEK inhibition in *NF1* mutant gliomas,^11,12^ and the MEK inhibitor selumetinib is FDA approved for tumors arising in patients with syndromic neurofibromatosis type 1 (NF-1) harboring a germline *NF1* mutation.^13,14^ In NF-1 associated gliomas, MEK inhibition demonstrates efficacy in a limited case series,^15^ and combined BRAF/MEK inhibition shows efficacy in BRAF p.V600E mutant gliomas,^16^ further supporting the translational potential of Ras/Raf/MEK/ERK blockade within genetically defined glioma subtypes. Nevertheless, treatment resistance to molecular monotherapy remains a challenge,^17–20^ and the mechanisms underlying MEK inhibitor resistance in *NF1* mutant glioma are unknown.

Here, we integrate targeted DNA sequencing, DNA methylation profiling, and single nuclear RNA-sequencing (snRNA-seq) of human patient somatic *NF1* mutant, IDH-wildtype glioblastomas with single cell RNA-sequencing (scRNA-seq), genome-wide clustered regularly interspaced short palindromic repeats interference (CRISPRi) screens, and pharmacologic studies in cell lines and mouse intracranial glioblastoma models to define molecular subgroups and functional mediators of MEK inhibitor response. Targeted DNA sequencing of *NF1* mutant, IDH-wildtype glioblastomas (n=186 tumors) revealed *CDKN2A/B* deletion was associated with poor outcomes in *NF1* mutant, but not *NF1* wildtype, glioblastomas. DNA methylation profiling (n=129 tumors) demonstrated three epigenetic subgroups with distinct clinical features, co-mutation patterns across cell cycle genes, and reference methylation classifier identities.

Genome-wide CRISPRi screens in mouse SB28 and human GBM43 glioblastoma cells identified a conserved cell cycle gene network mediating cell growth, consistent with the clinical importance of additional hits affecting the cell cycle in human somatic *NF1* mutant glioblastomas. Moreover, genome-wide mediators of selumetinib response converged upon two Ras pathway effectors mediating selumetinib sensitivity: *BRAF* and *SHOC2. SHOC2* repression in glioblastoma cells significantly improved selumetinib response both *in vitro* and in intracranial allografts *in vivo*. Single cell RNA-sequencing (scRNA-seq) of mouse intracranial glioblastomas treated with the MEK inhibitor selumetinib revealed MES-like tumor cells correlated with *CDKN2A* retention and the CRISPRi screen selumetinib sensitivity signature, with selumetinib resistant cells displaying Ras pathway induction. In contrast, non-MES like tumor cells were *CDKN2A* deficient and lacked expression of the CRISPRi screen selumetinib sensitivity signature, with selumetinib resistant cells inducing a glial de-differentiation program. Finally, snRNA-seq of *NF1* mutant, *CDKN2A/B* deleted, IDH-wildtype glioblastomas (n=9) showed non MES-like tumor cells exhibit increased cell cycle progression and were not associated with the CRISPRi screen selumetinib sensitivity signature. MES-like tumor cells within newly diagnosed, but not recurrent, tumors retained expression of the CRISPRi screen selumetinib sensitivity signature, suggesting resistance can arise both between and within specific transcriptomic glioblastoma cell tumor cell subpopulations. Taken together, our data identifies clinically important subgroups of *NF1* mutant, IDH-wildtype glioblastomas and supports a model in which heterogeneity between tumors and within tumor cell subpopulations underlies MEK inhibitor response, supporting the need for additional synergistic therapeutic approaches beyond maximal Ras pathway blockade for *NF1* mutant glioblastomas.

## Results

### DNA mutational analysis of somatic NF1 mutant, IDH-wildtype glioblastomas identifies CDKN2A/B loss as a negative prognostic factor

To define the genetic landscape of *NF1* mutant, IDH-wildtype glioblastoma, we retrospectively identified 186 newly diagnosed glioblastomas that underwent targeted DNA sequencing and harbored a somatic *NF1* mutation (Supplementary Table 1). Neuropathology review was performed to confirm that all cases met diagnostic criteria for this tumor type using the 2021 Central Nervous System Tumors WHO guidelines.^10^ Cases with a history of neurofibromatosis type 1, germline NF1 mutation, or methylation class match to high-grade astrocytoma with piloid features (HGAP) were excluded. Targeted sequencing across *NF1* mutant, IDH-wildtype glioblastomas identified recurrent alterations occurring in at least 10% of tumors including the *TERT* promoter, cell cycle genes (*CDKN2A/B, RB1, CDK4)*, phosphoinositide 3-kinase (PI3K) signaling (*PTEN, PIK3CA, PIK3R1)* or apoptosis (*TP53, MDM2, MDM4)* consistent with prior glioblastoma genomic analysis (Figure 1a).^1–3^ We next identified a propensity score matched *NF1* wildtype, IDH-wildtype glioblastoma cohort (Supplementary Table 2). When comparing patterns of mutational co-occurrence between *NF1* mutant and *NF1* wildtype glioblastomas, *NF1* mutant glioblastomas were significantly more likely to harbor co-mutation of *PTPN11*, an upstream Ras regulator, while *NF1* wildtype glioblastomas were significantly more likely to harbor *EGFR* alterations, and loss of the cell cycle/apoptosis genes *CDK4, MDM2,* or *MDM4* (Figure 1b). With regard to clinical outcomes, no significant difference in overall survival (OS) was observed between *NF1* mutant and *NF1* wildtype glioblastomas (Figure 1c). Cox proportional hazards (CPH) analysis of recurrently co-mutated genes occurring in at least 10% of the cohort revealed *CDKN2A/B* loss was associated with worse OS in *NF1* mutant but not *NF1* wildtype glioblastomas (Figure 1d-e; Supplementary Table 3). CPH analysis of clinical variables associated with OS in the *NF1* mutant, IDH-wildtype glioblastoma cohort revealed age, Karnofsky Performance Status (KPS),^32,33^ multifocality, adjuvant radiation,^34,35^ adjuvant temozolomide,^36,37^ and tumor treating fields (TTF)^38^ as being associated with OS (p<0.1). On multivariable CPH analysis incorporating significant clinical variables, *CDKN2A/B* loss remained significantly associated with worse OS in *NF1* mutant glioblastoma (Figure 1f; Supplementary Table 4). Taken together, DNA mutational analysis suggests *NF1* mutant and *NF1* wildtype tumors exhibit differences in co-mutation patterns with no global difference in clinical outcomes, and *CDKN2A/B* loss is a negative prognostic marker in *NF1* mutant but not *NF1* wildtype glioblastomas.

**Figure 1.**
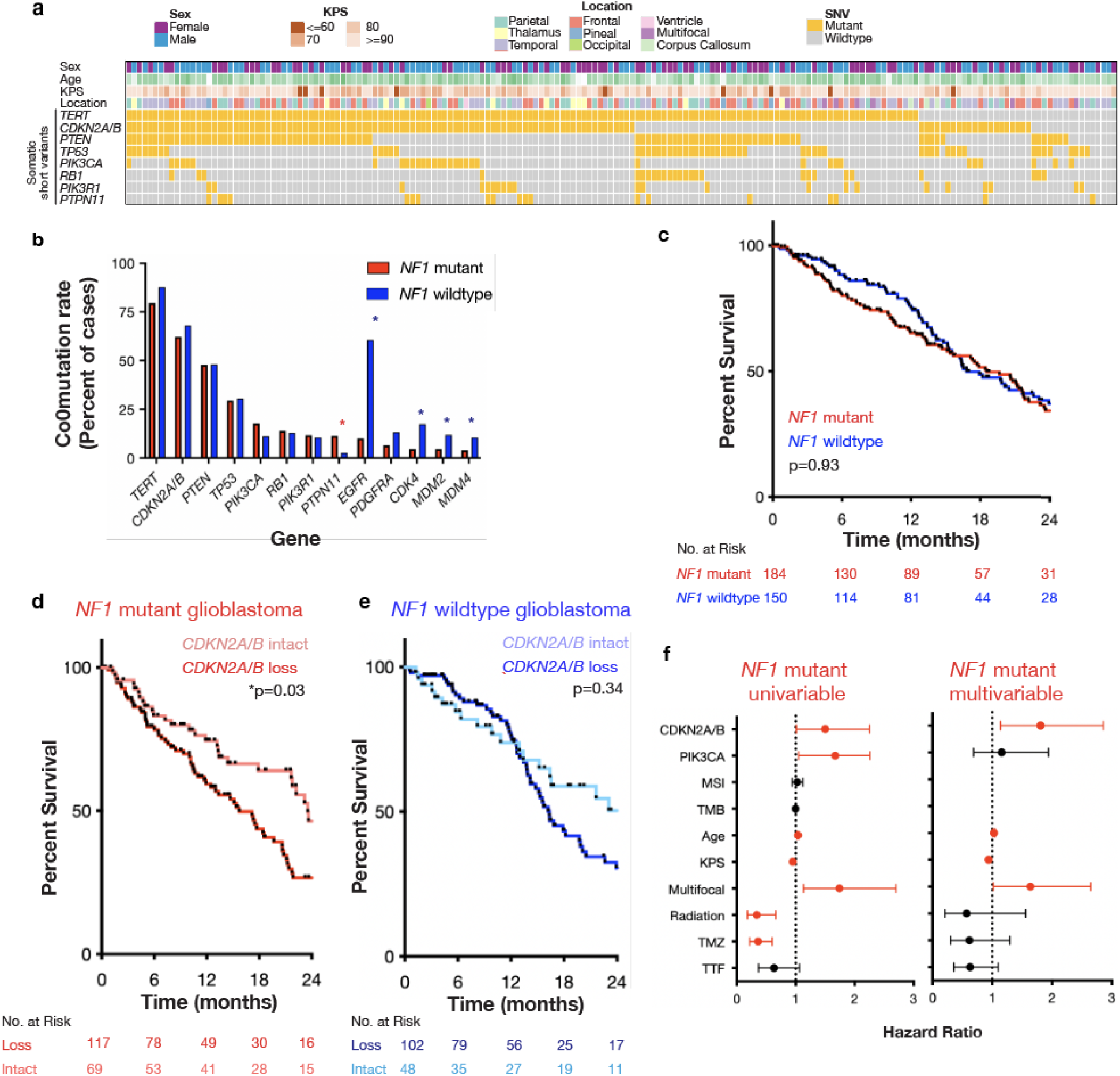
Targeted DNA sequencing of somatic *NF1* mutant, IDH-wildtype glioblastoma (n=186) reveals *CDKN2A/B* loss is associated with significantly worse overall survival. a. DNA mutational analysis of *NF1* mutant, IDH-wildtype glioblastoma (n=186) identifies recurrent co-alterations (>10% of tumors) in the *TERT* promoter*, TP53,* cell cycle regulators *(CDKN2A/B, RB1),* PI3K signaling (*PTEN, PIK3CA, PIK3R1)* and Ras signaling (*PTPN11)*. b. Analysis of recurrently co-mutated genes in *NF1* mutant and propensity score matched *NF1* wildtype glioblastoma cohort reveals *NF1* mutant tumors exhibit significantly increased co-alteration with *PTPN11* mutation and are mutually co-exclusive with *EGFR*, *CDK4, MDM2,* and *MDM4* alteration. c. *NF1* mutant glioblastomas do not demonstrate significantly different outcomes compared to *NF1* wildtype glioblastomas in a propensity score matched cohort of IDH-wildtype glioblastomas. d. *CDKN2A/B* loss is associated with significantly worse overall survival compared to *CDKN2A/B* intact tumors across *NF1* mutant but not e. *NF1* wildtype glioblastomas. f. Multivariable Cox proportional hazards analysis of recurrent genetic alterations shows *CDKN2A/B* loss is the only significant independent genetic factor associated with overall survival in *NF1* mutant glioblastoma.

### DNA methylation profiling reveals three epigenetic groups with distinct clinical and molecular features

To connect DNA mutations to epigenetic signatures,^9^ we next performed DNA methylation array profiling on *NF1* mutant, IDH-wildtype glioblastomas (n=129). Consensus unsupervised hierarchical clustering revealed 3 tumor groups: Group 1 Pediatric type high grade glioma, H3 and IDH-wildtype (n=12), Group 2 MES enriched (n=66), and Group 3 RTK enriched (n=51) (Figure 2a; Supplementary Figure 3a-b; Supplementary Table 5). No significant difference in overall survival was observed between methylation groups (Supplementary Figure 3c). With regard to baseline clinical parameters, significant differences were observed for age at diagnosis (Supplementary Figure 3d) and patient sex (Supplementary Figure 3e), with Group 1 tumors seen in younger patients, Group 2 tumors more likely to occur in male patients, and Group 3 tumors more likely to occur in female patients. No significant differences were observed between groups for KPS (Supplementary Figure 3f) or tumor location (Supplementary Figure 3g). Analysis of co-mutated genes by epigenetic group revealed Group 1 tumors were less likely to harbor *TERT* promoter alterations (p=0.005, Chi-square test) and more likely to harbor *TP53* mutations (p=0.03, Chi-square test), Group 2 tumors were more likely to contain *RB1* mutation (p=0.03, Chi-square test), and Group 3 tumors were more likely to exhibit *CDKN2A/B* homozygous deletion (p=0.001, Chi-square test) (Figure 2b). Classification using the Heidelberg random forest classifier (brain v12.8) revealed significant differences between groups (Figure 2c), with Group 1 tumors showing predominantly diffuse pediatric-type high grade glioma, H3- and IDH-wildtype, Group 2 tumors showing increased tumors with no match using the Heidelberg classifier, and Group 3 tumors enriched for glioblastoma IDH-wildtype receptor tyrosine kinase 1/2 (RTK1/2) tumors. Methylation classification was associated with significant differences in OS (Figure 2d); MES and adult type diffuse gliomas showing poor outcomes and tumors with no classification showing the best outcomes, with RTK tumors demonstrating an intermediate clinical course. In sum, DNA methylation profiling revealed 3 epigenetic groups of somatic *NF1* mutant, IDH-wildtype glioblastomas associated with distinct clinical features, co-mutation in patterns in cell cycle genes such as *TP53, RB1,* and *CDKN2A/B*, and reference methylation classification assignments, underscoring the heterogeneity between these tumors.

**Figure 2.**
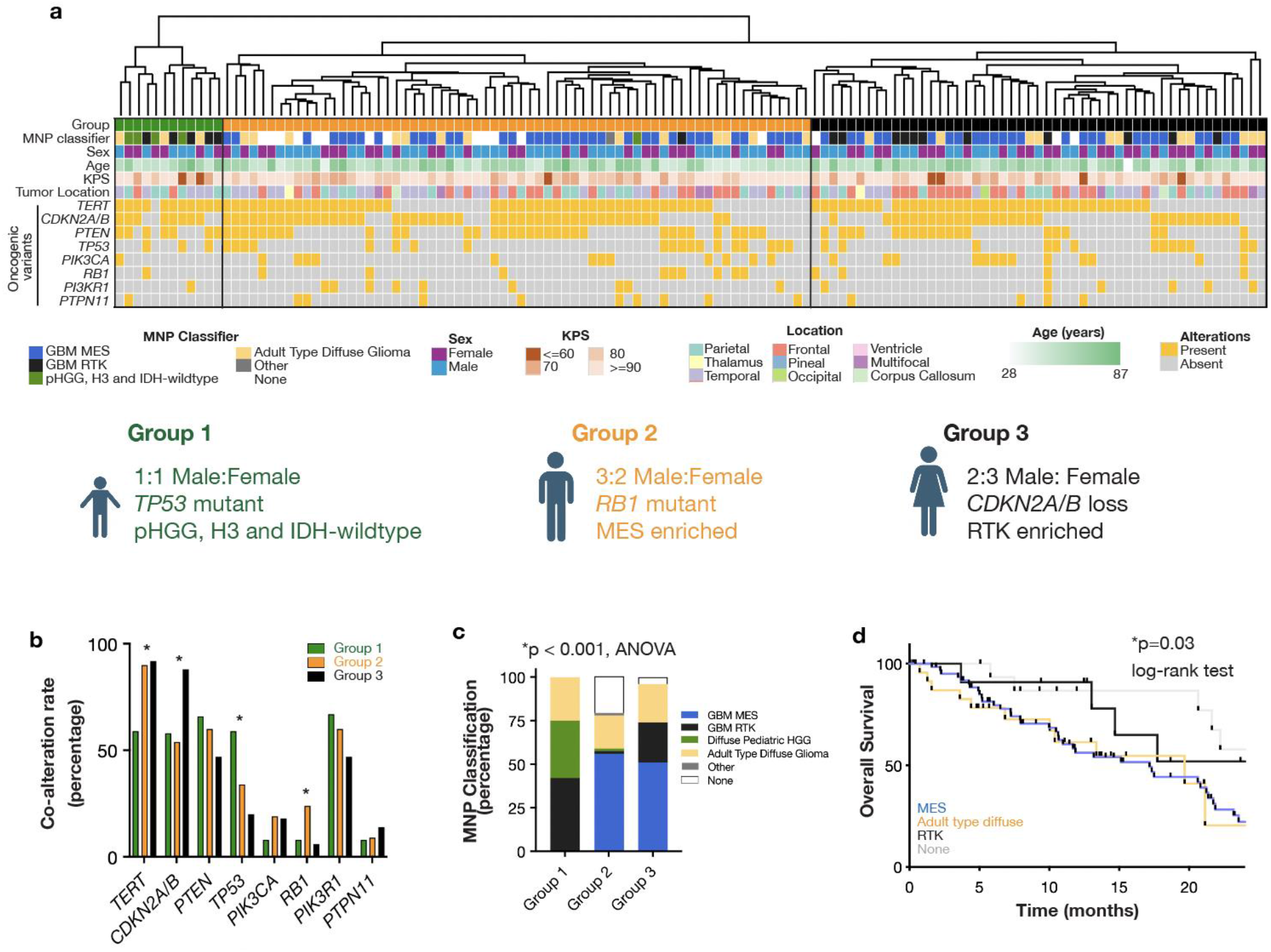
DNA methylation profiling reveals three epigenetic groups of *NF1* mutant, IDH*-* wildtype glioblastomas with distinct clinical and molecular features. a. DNA methylation array profiling (n=129) of *NF1* mutant, IDH-wildtype glioblastoma reveals three epigenetic groups. b. *NF1* mutant glioblastoma DNA methylation group is not associated with significant differences in overall survival (p=0.35, log-rank test). DNA methylation groups demonstrate significant differences in c. co-alteration rates in *TERT* mutation (p=0.005, Chi-square test), *CDKN2A/B* loss (p=0.001, Chi-square test), *TP53* mutation (p=0.03, Chi-square test), and *RB1* mutation (p=0.03, Chi-square test), c. molecular neuropathology (MNP) methylation classification (p<0.0001, Chi-square test) and d. are associated with overall survival (p=0.03, log-rank test) by MNP classification in *NF1* mutant glioblastomas.

### Genome-wide CRISPRi screens in glioblastoma cells demonstrate conserved mechanisms underlying cell growth and response to the MEK inhibitor selumetinib

To more broadly define genetic perturbations important for *NF1* mutant glioblastoma cell growth and responses to MEK inhibition, a therapeutic strategy for *NF1* mutant tumors, we performed genome-wide CRISPRi screens in two cell models of *NF1* mutant glioblastoma: *NF1* mutant human GBM43 cells and *Nras^G12V^* mutant mouse SB28 cells (Figure 3a; Supplementary Table 6 and 7). We first compared T10 DMSO treated cells to T0 cells to identify genes required for glioblastoma cell growth. In SB28 cells, a total of 1,445 genes led to significantly decreased cell growth while 137 genes led to increased cell growth, with *Cdkn2a* and *Tp53* loss comprising two of the top hits required for cell growth consistent with human tumor mutation data (Supplementary Figure 4a). In GBM43 cells, a total of 690 genes led to significantly decreased cell growth while 61 genes led to increased cell growth (Supplementary Figure 4b), and gene ontology analysis of genes required for cell growth in either cell line converged on regulators of cell cycle progression (Supplementary Figure 4c-d). By integrating genes required for growth in both screens, a 31 gene consensus cell cycle signature required for *NF1* mutant glioblastoma cell growth was identified, enriched for components within the cyclin dependent kinase (CDK) / retinoblastoma (RB) or TP53 (Figure 3b) co-mutations observed in human tumors (Figure 1a).

**Figure 3.**
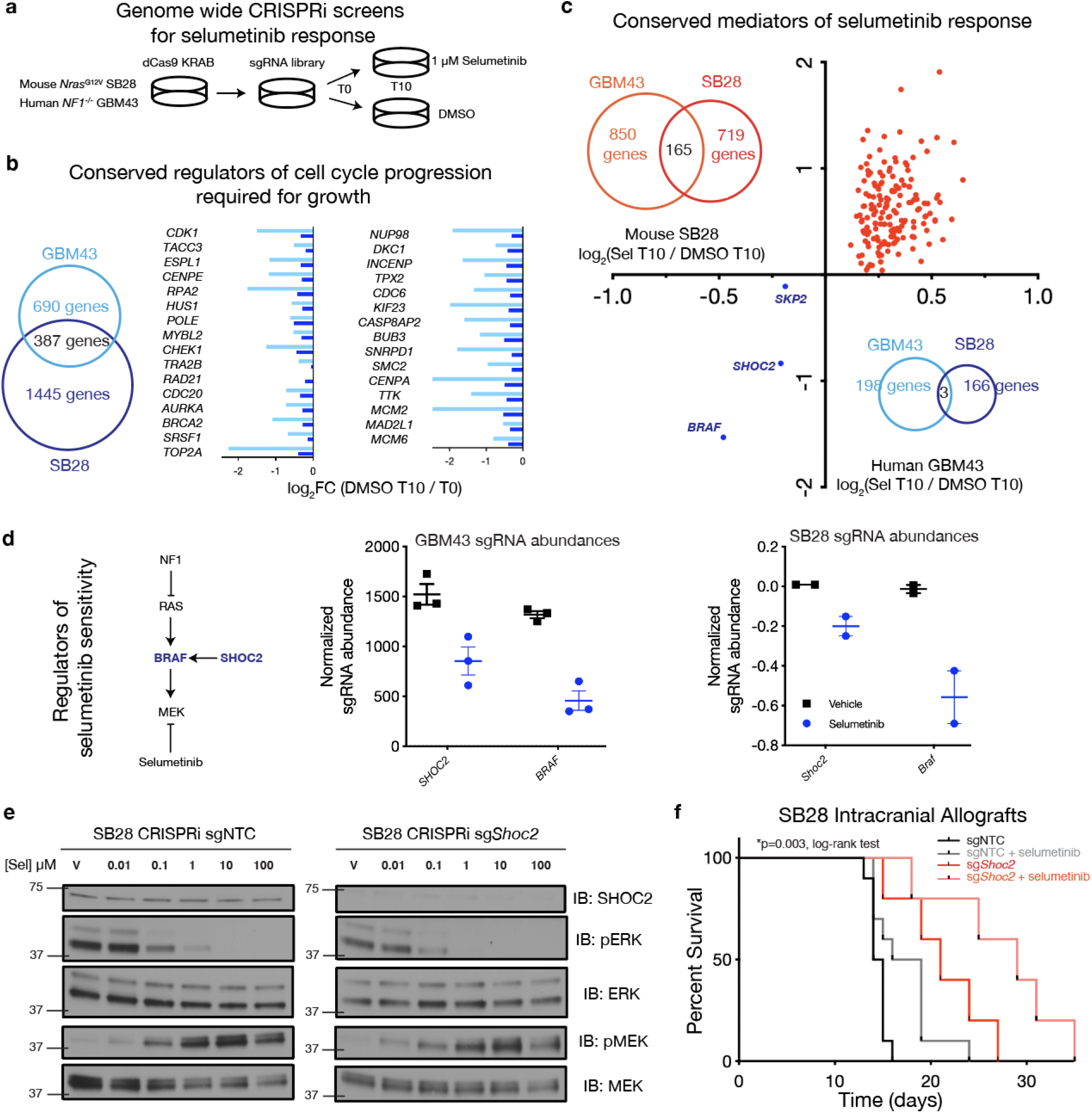
Genome-wide CRISPRi screens in glioblastoma cells reveal cell cycle genes are required for growth and nominates *SHOC2* as a critical mediator of MEK inhibitor selumetinib. a. Schematic of genome-wide CRISPRi screens in *NF1* mutant human GBM43 (n=3) and *NRAS^G12V^* mouse SB28 (n=2) glioblastoma cells treated with 1 μM selumetinib or vehicle DMSO. b. Integrated analysis of genes significantly mediating cell growth (DMSO / T0) reveals a 31 gene conserved cell cycle network required for SB28 and GBM43 cell growth. c. Integrated analysis of conserved genes significantly modulating glioblastoma cell response to the MEK inhibitor selumetinib (T10 Selumetinib / T10 DMSO) in both SB28 and GBM43 screens (n=165 genes mediating resistance, n=3 genes mediating sensitivity) reveals d. repression of the Ras downstream targets *BRAF* or *SHOC2* mediates selumetinib sensitivity. e. sg*Shoc2* deficient SB28 glioblastoma cells (IC50 0.0751, 95% CI 0.011 – 0.027) are more sensitive to selumetinib than sgNTC SB28 glioblastoma cells (IC50 0.1131, 95% CI 0.080 – 0.169) (p<0.0001, F-test). f. CRISPRi *Shoc2* repression is sufficient to improve selumetinib response in SB28 intracranial allografts *in vivo*.

We next evaluated genetic perturbations selectively altering selumetinib response by comparing sgRNA abundance between cells treated with selumetinib for 10 days (T10 Selumetinib) and compared to cells treated with vehicle DMSO for 10 days (DMSO T10). In SB28 cells, a total of 166 genes mediated sensitivity while 719 genes mediated resistance while in GBM43 cells, while 198 genes mediated sensitivity and 850 genes mediated resistance (Supplementary Figure 5a-b); integration of conserved hits across both screens identified a total of 3 genes mediating sensitivity and 165 genes mediating resistance (Figure 3c). We focused on genes mediating selumetinib sensitivity as potentially druggable dependencies, and STRING based protein network analysis revealed highly connected enrichment of Ras signaling nodes amongst sensitizing hits in both SB28 and GBM43 cells (Supplementary Figure 5c-d). Indeed, the top 2 conserved sensitivity hits were *BRAF,* a direct effector of RAS with established roles in glioma,^16,17,39^ and *SHOC2,* which has not been previously implicated in glioblastoma (Figure 3d). SHOC2 functions as part of a heterotrimeric complex to remove an inhibitory phosphorylation on RAF and thus promote RAF activation of MEK.^40,41^ Given its role as a potentially novel therapeutic target in glioblastoma, we next validated SHOC2 as a mediator of selumetinib response. *SHOC2* repression in SB28 or GBM43 glioblastoma cells (Supplementary Figure 6a-b) followed by biochemical analysis of selumetinib response revealed CRISPRi *Shoc2* deficient SB28 cells (Figure 3e; Supplementary Figure 6c) and siRNA *SHOC2* deficient GBM43 cells (Supplementary Figure 6d) exhibited increased sensitivity to selumetinib. We next tested the effect of *Shoc2* repression and selumetinib treatment given at a dose of 25 mg/kg twice daily in SB28 intracranial tumors, demonstrating that *Shoc2* loss combined with selumetinib resulted in more durable responses compared to either perturbation alone (Figure 3f; Supplementary Figure 7). Taken together, genome-wide CRISPRi screens suggest cell cycle dysregulation is essential for *NF1* mutant glioblastoma growth and compensatory Ras activation underlies MEK inhibitor responses, with *SHOC2* constituting a potential additional therapeutic target to maximally block Ras pathway output and enhance selumetinib responses in glioblastoma.

### Different tumor cell subpopulations exhibit distinct transcriptional signatures underlying selumetinib response in SB28 glioblastoma intracranial tumor models

Although combined *Shoc2* repression and selumetinib significantly improved responses in SB28 intracranial glioblastomas, tumors ultimately developed resistance and the therapeutic effect was not durable. To define the transcriptomic and cellular mechanisms underlying selumetinib resistance in SB28 intracranial tumors, we next performed scRNA-seq from SB28 intracranial tumors treated with either vehicle (n=3) or selumetinib (n=3) (Figure 4a; Supplementary Figure 8a-c). Non-tumor cells were identified and filtered out based on *Ptprc* expression and cell type classification (Supplementary Figure 8d-f), retaining a total of 3,411 tumor cells distributed across 6 tumor cell clusters in Uniform Manifold Approximation and Projection (UMAP) analysis (Figure 4b; Supplementary Figure 9a-b; Supplementary Table 8). Tumor cell transcriptional states were assigned based on published glioblastoma cell state signatures,^7^ identifying 3 MES-like clusters, 1 NPC-like cluster, 1-OPC-like cluster, and 1 AC-like cluster, and tumor cell type composition was not significantly different based on treatment condition (Figure 4c; Supplementary Figure 9c-d). MES-like tumor clusters retained *Cdkn2a* expression while non-MES like tumor cells lacked *Cdkn2a* expression (Figure 4d; Supplementary Figure 9e). To connect transcriptomic signatures to functional mediators of selumetinib response, we constructed a 165 gene signature comprising the conserved hits mediating selumetinib resistance in both CRISPRi screens as a selumetinib sensitivity signature. We found that MES-like cells were associated with a significantly increased selumetinib sensitivity signature compared to non MES-like cells (Figure 4e; Supplementary Figure 9f). Finally, to understand mechanisms associated with selumetinib resistance, we performed differential expression analysis between selumetinib treated compared to vehicle treated tumor cell populations (Supplementary Table 9). While selumetinib treated MES-like cells exhibited Ras pathway activation evidenced by significantly increased expression of the Ras transcriptional targets *Fos, Jun, Junb, Jund,* and *Dusp1* (Figure 4f), non MES-like cells treated with selumetinib instead were associated with induction of glial de-differentiation genes such as *Olig1, Olig2, Sox10, Ptprz1,* and *Pdgfra,*(Figure 4g). In sum, these data suggest distinct mechanisms mediate selumetinib response across different glioblastoma cellular states, with MES-like cells maintaining dependence on Ras/Raf/MEK/ERK signaling while non MES-like cells activate a de-differentiation program to persist following selumetinib treatment.

**Figure 4.**
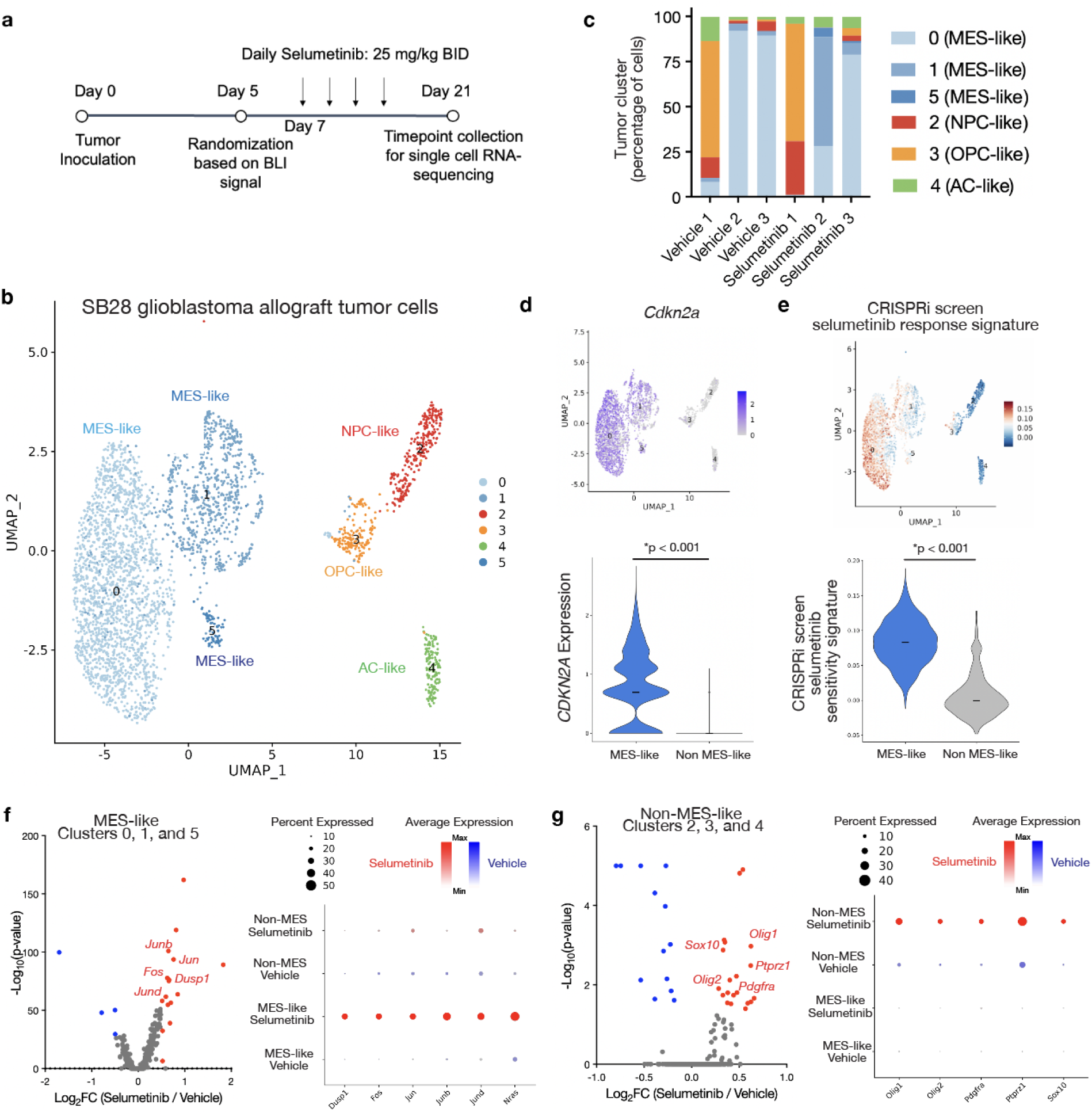
Single cell RNA-sequencing of SB28 intracranial allografts reveals distinct transcriptional signatures and selumetinib responses in MES-like compared to non MES-like tumor cell subpopulations. a. Experimental overview of single cell RNA-sequencing of selumetinib versus vehicle treated SB28 intracranial allografts. b. Uniform Manifold Approximation and Projection (UMAP) analysis of 3,411 glioblastoma SB28 intracranial allograft tumor cells reveals 6 tumor cell clusters corresponding to 3 MES-like subpopulations, 1 NPC-like subpopulation, 1 OPC-like subpopulation, and 1 AC-like subpopulation. c. Breakdown of tumor cell subpopulations by treatment condition reveals no significant differences in cell type composition between selumetinib and vehicle treated tumors. d. MES-like tumor cells retain *Cdkn2a* expression while non MES-like tumor cells lack *Cdkn2a* expression. e. Integration of a consensus CRISPRi screen selumetinib response signature (n=165 conserved genes mediating resistance across both SB28 and GBM43 screens) shows MES-like cells are associated with selumetinib response. f. Selumetinib treated MESl-like cells show significantly increased expression of Ras pathway transcriptional targets such as *Fos, Jun, Junb, Jund,* and *Dusp1.* g. Selumetinib treated non- MES-like cells show significantly increased expression of de- differentiation genes such as *Olig1, Olig2, Sox10, Ptprz1,* and *Pdgfra*.

### Single nuclear RNA-sequencing of human NF1 mutant glioblastoma reveals MES-like cell state is associated with selumetinib sensitivity

To investigate the intra-tumor heterogeneity across somatic *NF1* mutant, *CDKN2A/B* homozygous deleted, IDH-wildtype glioblastomas and how this relates to our observations regarding selumetinib responses in glioblastoma models, we next performed snRNA-seq on a total of 21,959 nuclei from 9 human patient-derived glioblastomas (Figure 5a). Integrated UMAP revealed a total of 14 cell clusters comprising 5 tumor cell clusters and 9 non-tumor cell clusters defined by *PTPRC (*CD45), marker gene expression, and CNAs (Figure 5a; Supplementary Figure 10a-e; Supplementary Table 10).^42,43^ Tumor cells showed recurrent chromosome 7 gain, chromosome 10 loss, chromosome 13 loss, or chromosome 19 gain (Supplementary Figure 10e). Non-tumor cell clusters included oligodendrocytes (C2), microglial cells (C4, C7, C12), neurons (C8, C9, C10), T cells (C11), and endothelial cells (C13) (Supplementary Figure 11a-b). Using published glioblastoma cell state signatures,^7^ tumor clusters were classified as MES-like (C1, C5), AC-like (C3), NPC-like (C6), or OPC-like (C0) (Figure 5a; Supplementary Figure 11c). Assignment of tumor transcriptional states by sample revealed 6 out of 9 samples (S1-6) were enriched for MES-like clusters, although all 9 tumors contained multiple tumor cell subpopulations consistent with an admixture of tumor cell states comprising each individual tumor (Figure 3b). Cell cycle signature scoring showed that MES-like tumor cells contained a significantly decreased number of cycling cells (Figure 3c) with significantly decreased S phase cells compared to non MES-like tumor cells (Figure 3d). The CRISPRi screen selumetinib sensitivity signature was significantly increased in MES-like compared to non MES-like cells (Figure 3e), but this increase was limited to MES-like cells in the primary, but not recurrent, tumor samples (Figure 3f). In sum, snRNA-seq of human *NF1* mutant, *CDKN2A/B* deleted glioblastomas supports a model in which selumetinib sensitivity is restricted to MES-like cells while non MES-like cells exhibit increased cell cycle progression through parallel mechanisms independent of Ras/Raf/MEK/ERK signaling uncoupled from selumetinib response.

**Figure 5.**
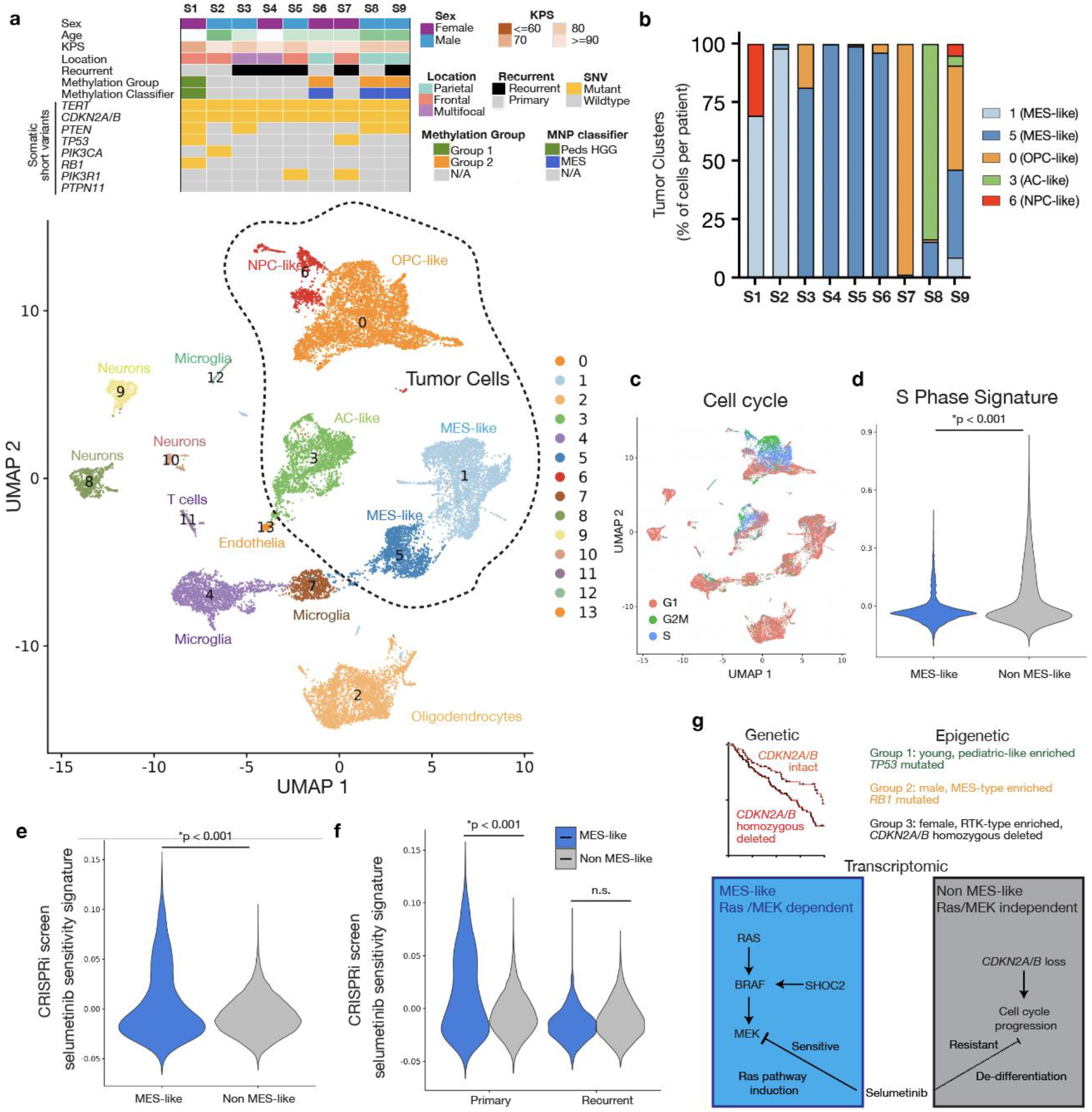
Single nuclear RNA-sequencing of human *NF1* mutant, IDH-wildtype glioblastoma reveals MES-like cells in the primary, but not recurrent, tumor are associated with selumetinib sensitivity. a. Single nuclear RNA-sequencing (snRNA-seq) of 21,959 nuclei from 9 patient derived human *NF1* mutant, *CDKN2A/B* homozygous deleted, IDH- wildtype glioblastomas identifies 5 tumor cell clusters and 9 non-tumor cell clusters. b. Assignment of published glioblastoma cellular states by sample of origin reveals both inter and intra-tumor heterogeneity, with 6 of 9 *NF1* mutant glioblastomas predominantly comprised of MES-like clusters. c. Cell cycle assignment shows MES-like tumor cell clusters are associated with decreased cycling cells in d. S phase compared to non MES-like cells. e. The CRISPRi screen selumetinib response signature is significantly enriched in MES-like tumor cells compared to non MES-like cells (p<0.001, Wilcoxon test). f. MES-like cells in recurrent tumors no longer express the CRISPRi screen selumetinib sensitivity gene signature (p<0.001, Wilcoxon test). g. Genetic, epigenetic, and transcriptomic heterogeneity underlie clinical differences and selumetinib response in *NF1* mutant glioblastomas.

## Discussion

*NF1* loss is recurrently mutated in glioblastoma yet the efficacy of targeted therapies leveraging this genetic alteration is limited. Here, we combine multiplatform bulk and single cell genomic analysis of somatic *NF1* mutant, IDH-wildtype glioblastomas with genome-wide CRISPRi screens and mouse intracranial tumor models to better understand the molecular landscape and MEK inhibitor responses in these tumors. Targeted DNA sequencing revealed *EGFR* alteration is mutually exclusive with *NF1* mutation as previously reported,^1–3^ and moreover, *CDKN2A/B* homozygous deletion in *NF1* mutant, but not *NF1* wildtype, glioblastoma is associated with worse survival. DNA methylation profiling identified three epigenetic groups with differences in age, biologic sex, co-mutated tumor suppressors, and methylation classification, and methylation classifier groups exhibited significant differences in clinical outcome, further underscoring the heterogeneity across *NF1* mutant, IDH-wildtype glioblastomas. Genome-wide CRISPRi screens in two different glioblastoma cell lines demonstrated additional perturbations in cell cycle genes are required for cell growth while selumetinib sensitivity is dependent on additional Ras pathway outputs, identifying *Shoc2* loss as sufficient to improve selumetinib responses in mouse intracranial glioblastoma tumors.

However, combined *Shoc2* repression plus selumetinib treatment did not result in a durable long-term response, and scRNA-sequencing of vehicle or selumetinib treated mouse intracranial tumors demonstrated differences in *CDKN2A* expression, selumetinib sensitivity, and transcriptional programs underlying selumetinib resistance between MES-like and non MES-like tumor cell subpopulations. Consistent with these observations from genome-wide CRISPRi screens and mouse intracranial tumors, snRNA-seq of human *NF1* mutant, *CDKN2A/B* deleted glioblastomas revealed multiple transcriptional subtypes within and between patients, linking MES-like cell states to decreased cell cycle progression and increased selumetinib sensitivity.

More broadly, it appears the MES-like transcriptional signature classically associated with *NF1* mutation can be effectively treated with maximal Ras/Raf/MEK/ERK pathway blockade while non MES-like cells leverage divergent cell cycle and de-differentiation mechanisms to resist Ras/Raf/MEK/ERK pathway blockade. Taken together, our data support a model in which heterogeneity exists at the genetic, epigenetic, and transcriptomic levels that underlie clinical outcomes and treatment response within *NF1* mutant glioblastomas (Figure 5g).

The efficacy of molecular monotherapy for malignant tumors such as glioblastoma is limited by numerous factors including tumor heterogeneity, compensatory intracellular signaling changes, and cellular plasticity, motivating approaches to understand, and ultimately circumvent, treatment resistance. Based on our data, devising novel approaches to fully block Ras pathway outputs such as through *SHOC2* perturbation or combined pan-RAF^39^ plus MEK inhibition may show benefit in depleting the subpopulation of MES-like tumor cells and improving clinical responses to targeted therapy. While no clinical compounds blocking SHOC2 exist to date, the recent structures of the SHOC2 heterotrimeric complex have facilitated potential pharmacologic strategies that may be useful in glioblastoma.^44–46^ However, synergistic combinations targeting the parallel susceptibilities of non MES-like cells will also be critical, and it will be important to define whether CDK4/6 inhibitors targeting *CDKN2A/B* loss may be useful in this context. The observed inter-tumor heterogeneity at the level of DNA alterations and methylation as well as intra-tumor heterogeneity observed in mouse and human single cell RNA-sequencing data^7,47^ underscores the importance of patient selection within *NF1* mutant glioblastomas. In that regard, the identification of a functional genomic selumetinib sensitivity signature based on our CRISPRi screens may provide a useful tool to identify transcriptional subpopulations, and the patients with tumors that harbor them, in which Ras pathway blockade will prove effective.

In sum, our data support the existence of clinically significant inter-tumor and intra-tumor differences within *NF1* mutant, IDH-wildtype glioblastoma with distinct responses to the MEK inhibitor selumetinib. Future work leveraging larger, multi-institutional cohorts will be critical to validate the observation that *CDKN2A/B* loss is a negative prognostic marker and the epigenetic groups observed in the present work, all of which were obtained from samples at a single institution. Similarly, the relationship between putative glioblastoma mutational group, single cell composition, and transitions between primary and recurrent tumors following different therapies will require further investigation to define *NF1* specific cellular mechanisms underlying targeted therapy resistance. Finally, future preclinical and clinical trials aimed at complementing MEK inhibitors with alternate modalities such as radiation, immunotherapy, or targeted agents against co-mutated genes such as *CDKN2A/B* will be critical to achieve durable responses in *NF1* mutant glioblastomas.

## Acknowledgments

We are grateful to Anny Shai and the staff of the UCSF Brain Tumor Center Biospecimen and Pathology Core for histological staining and tissue acquisition, Eric Chow and the staff of the UCSF Center for Advanced Technology for sequencing, and to the UCSF Brain Tumor Center Preclinical Therapeutics Core for assistance with intracranial tumor experiments. The GBM43 *NF1* mutant glioblastoma model was a gift from Jann Sakaria and the Mayo Clinic Brain Tumor Patient-Derived Xenograft National Resource. We thank CJ Lucas, Karisa Schreck, Matthew Sale, Rony Francois, Lucy Young, and Frank McCormick for helpful discussion and critical comments on this manuscript. Resources were provided by the UCSF Brain Tumor SPORE Biorepository (NIH 5P50 CA097257) and the Panattoni Family Research Program. This work was supported by a Neurofibromatosis Therapeutic Acceleration Program (NTAP) Francis Collins Scholar Award, a Children’s Tumor Foundation Young Investigator Award, UCSF Brain Tumor Center SPORE Career Enhancement Program (CEP) award, and a UCSF Helen Diller Family Comprehensive Cancer Center Neuro-Oncology Research Grant to H.N.V.

## Online Methods

### Clinical database design and DNA sequencing of human glioblastomas

Patients treated with surgical resection for a glioblastoma at the University of California San Francisco from 2006-2024 were retrospectively identified from a tissue biorepository. Male and female patients over 18 years of age were included. A total of 186 newly diagnosed IDH-wildtype glioblastoma cases containing a pathogenic or likely pathogenic somatic *NF1* mutation using a clinical CLIA certified targeted DNA next generation sequencing assay obtained as part of routine clinical care were retrospectively identified. Cases were verified to meet histologic and molecular diagnostic criteria for IDH-wildtype glioblastoma as specified in the 2021 WHO classification of central nervous system tumors. Exclusion criteria included any patients with a clinical history or diagnosis of neurofibromatosis type 1, a germline *NF1* mutation, or tumors that matched to high-grade astrocytoma with piloid features (HGAP) using v12.8 of the Heidelberg DNA methylation classifier. Demographic and clinical information was extracted from the electronic medical record and an institutional cancer registry. Survival status was obtained through searching the electronic medical record, institutional cancer registry, Social Security, Department of Motor Vehicles (DMV), nationwide hospital databases, and publicly available obituaries. A summary of demographic information is included in Table 1. This study was approved by the UCSF Institutional Review Board, protocol 10-01318 and 22-37134.

**Table 1.** Clinical parameters and molecular analysis within the single institution cohort of newly diagnosed *NF1* mutant, IDH-wildtype glioblastomas.

| Parameter | Value (n=186 tumors) |
| --- | --- |
| **Sex** | n=98 male (53%)<br>n=88 female (47%) |
| **Age (median; range)** | 60.47 years (20.47-86.96) |
| **Median Survival** | 11.7 months |
| **KPS** |  |
| <=60 | 9 (4%) |
| 70 | 27 (15%) |
| 80 | 59 (32%) |
| >=90 | 90 (49%) |
| **Tumor location** |  |
| Parietal | 42 (23%) |
| Frontal | 57 (30%) |
| Occipital | 1 (1%) |
| Temporal | 60 (32%) |
| Thalamus | 11 (6%) |
| Corpus Callosum | 3 (2%) |
| Pineal | 3 (2%) |
| Ventricle | 1 (1%) |
| Multifocal | 7 (4%) |
| **DNA Methylation Profiling** | n=129 (70%) |
| **Postoperative radiation** | n=164 (89%) |
| **Postoperative temozolomide** | n=148 (80%) |
| **Tumor treating fields** | n=32 (17%) |

### DNA Methylation Profiling

Methylation profiling was performed using the Illumina EPIC (V1: n=47; V2: n=82) platform on sporadic *NF1* mutant IDH-wildtype glioblastomas. Archival FFPE tissue blocks were screened for areas with high tumor content. 2.0 mm tissue punches were used to extract FFPE tissue from blocks with adequate material. DNA was extracted using the Qiagen FFPEasy kit and 1500 ng DNA was loaded per sample. DNA concentrations down to a minimum of 20ng/µl (total: 1.5µg) were considered acceptable for methylation array profiling. DNA was processed according to manufacturer’s using the Infinium FFPE QC and Restore Kit (WG-321-1001 and WG-321-1002, Illumina) followed by the EPIC beadchip (WG-317–1003, Illumina). All downstream analysis were performed in R using the minfi Bioconductor package as previously reported.^20–23^ Data were normalized via functional normalization and filtered based on the following criteria: (i) removal of probes targeting the X and Y chromosomes, (ii) removal of probes containing a common single nucleotide polymorphism (SNP) within the targeted CpG site or on an adjacent basepair, and (iii) removal of probes not mapping uniquely to the human reference genome. Unsupervised hierarchical clustering (Euclidean distance, complete linkage method) was performed using the top 5,000 most variable probes as previously described.^15^ β values were used for visualization of methylation levels (β=methylated/[methylated+unmethylated]) and M values were used for statistical analysis (M=log2[methylated/unmethylated]). ConsensusClusterPlus (v 1.62.0) analysis (hierarchical clustering, k range 2 to 10, 1000 repetitions) was used to assess optimal cluster size and stability.

### Human glioblastoma single nuclear RNA-sequencing (snRNA-seq)

Single nuclei were isolated from frozen human sporadic *NF1* mutant, IDH-wildtype glioblastoma surgical resection samples with greater than 30% tumor content using an automated tissue dissociation system (Singulator S100, S2 genomics, CA, USA). Briefly, 10 - 20µg of frozen tumor-rich sample was homogenized and purified using a nuclei isolation kit, buffer containing RNAase, and an NIC+ cartridge. The resulting suspension was purified using a 40 µm filter and a Percoll gradient. Nuclei were counted using an automated cell counter as well as DAPI staining. Single-nuclei suspensions were processed for single-nuclei RNA sequencing using the Chromium Single Cell 3’ GEM, Library & Gel Bead Kit v3.1 (1000121, 10x Genomics) and a 10x Chromium X controller, with cell suspensions diluted per the manufacturer recommended protocol and settings for a target recovery of 8,000 cells per sample. Libraries were sequenced on an Illumina NovaSeq 6000, targeting >50,000 reads per cell, at the UCSF Center for Advanced Technology. Library demultiplexing, read alignment, identification of empty droplets, and UMI quantification were performed using CellRanger (https://github.com/10xGenomics/cellranger).

All downstream analyses were performed with Seurat v4.3.0.^24^ Cellranger generated filtered feature matrices were imported into a Seurat object. For quality control, data were filtered on a per sample basis to remove outliers in gene count, UMI count, mitochondrial genes, and ribosomal genes. The individual count matrices were normalized by SCTransform v2.

Scanorama (https://github.com/brianhie/scanorama)^25^ was used to perform data integration across datasets and cluster number optimization was performed by comparing multiple cluster resolutions using Clustree and the selected cluster resolution was examined by silhouette-width analysis, which reported a mean width per cluster larger than 0. Genes differentially expressed in each cluster were identified using FindAllMarkers function (cutoff: min.pct = 0.25, logfc.threshold = 0.25, min.diff.pct = 0.2). Tumor versus non-tumor cell designation was performed through a combination of manual marker gene inspection and inferCNV (https://github.com/broadinstitute/inferCNV)^26^ of the CD45-negative clusters using endothelial cells as reference. SCEVAN^27^ (https://github.com/AntonioDeFalco/SCEVAN) was used as attempt to delineate tumor from non-tumor cells based on individual sample’s copy number alterations (CNAs). Cell type designation of non-tumor cells was performed through scType automated cell type classification.^28^ The automated cell type classification algorithms were developed for normal cells with clearly defined marker genes and become limited when applied to tumor ^27^cell types due to the ambiguity of gene signatures of malignant cell types. Therefore, alternative methods were explored to define the different cell states of tumor cells. Tumor subtypes were defined by gene signatures scoring of MES-like, AC-like, NPC-like and OPC-like cells as previously described.^7^ Signature scores of cells were performed with AddModuleScore function. Cell type classification was validated through a combination of manual marker gene inspection, gene ontology analysis, and cell cycle phase classification. ANOVA or Wilcoxon tests were used for multiple comparisons and Student’s t-test was used for two group comparisons.

### Mouse intracranial tumor establishment, bioluminescence imaging, and drug treatment

Five to six-week-old female C57BL/6 (Envigo Laboratories, Livermore, CA), housed under aseptic conditions, received intracranial tumor cell injection through the UCSF Brain Tumor Center Preclinical Therapeutics Core, and as approved by the University of California San Francisco Institutional Animal Care and Use Committee. Briefly, mice were anesthetized by combination of intraperitoneal injection of a mixture containing ketamine (100mg/kg) and xylazine (10 mg/kg), and inhalation of isoflurane. The scalp was surgically prepped, and a skin incision ∼10 mm in length was made over the middle frontal to parietal bone. The surface of the skull was exposed so that a small hole was made 3.0mm to the right of the bregma and just in front of the coronal suture with a 25-gauge needle. A 26-gauge needle attached to a Hamilton syringe was inserted into the hole in the skull. The needle was covered with a sleeve that limits the injection depth to 3-4mm. 3 μL of tumor cell suspension was injected into the right caudate putamen at a rate of 1 μL/min by free hand. The skull surface was then swabbed with hydrogen peroxide before the hole was sealed with bone wax to prevent reflux. The scalp was closed with surgical staple. Mice were treated with either selumetinib 25 mg/kg or with vehicle control by oral gavage twice daily. For bioluminescence imaging (BLI), mice were anesthetized with inhalation of isoflurane, then administered 150 mg/kg of luciferin (D-luciferin potassium salt, Gold Biotechnology, St. Louis, MO) via intraperitoneal injection. Ten minutes after luciferin injection, mice were examined for tumor bioluminescence with an IVIS Lumina imaging station and Living Image software (Caliper Life Sciences, Alameda, CA), and intracranial regions of interest were recorded as photons per second per steradian per square cm.

### Mouse intracranial allograft single cell RNA-sequencing

Harvested tumors were mechanically minced with forceps and then dissociated to single cell suspension using the Papain Dissociation System (Worthington #LK003150) following the manufacturer’s protocol. To further remove non-cellular debris, cell suspensions were passed through a 70 μM strainer (Corning, #352350), centrifuged at 300g for 5 minutes, and resuspended in cold phosphate buffered saline.

Similar to human single nuclear RNA-sequencing, intracranial allograft single-cell sequencing was performed using the Chromium Single Cell 3’ Library & Gel Bead Kit v3.1 on a 10X Chromium controller (10X Genomics) using the manufacturer recommended default protocol and settings. Samples were sequenced on an Illumina NovaSeq at the UCSF Center for Advanced Technology, and the demultiplexed FASTQ files were processed using CellRanger for alignment to the mm10 reference genome, identification of empty droplets, and determination of a count threshold. All downstream analyses were performed with Seurat v4.4.^24^ Cellranger generated filtered feature matrices were imported into a Seurat object. For quality control, data were filtered on a per sample basis to remove outliers in gene count, UMI count, mitochondrial genes, and ribosomal genes. The individual count matrices were normalized by SCTransform v2. Scanorama was used to perform data integration across datasets and cluster number optimization was performed by comparing multiple cluster resolutions using Clustree and the selected cluster number was verified with silhouette scores. Genes differentially expressed in each cluster were identified using FindAllMarkers function (cutoff: min.pct = 0.25, logfc.threshold = 0.25, min.diff.pct = 0.1). Tumor vs non-tumor cell designation was based on gene expression but not CNAs because the algorithms available for inferring CNAs were developed for human cells only. Non-tumor cells were identified by a combination of CD45 expression, manual marker gene inspection and scType automated cell type classification.

Tumor cells were then re-clustered and tumor subtypes were defined by gene signatures scoring of MES-like, AC-like, NPC-like and OPC-like cells as previously described.^7^ Genes differentially expressed in each cluster in selumetinib-treated compared to vehicle-treated samples were identified using FindMarkers function (cutoff: min.pct=0.2, logfc.threshold=0) and the results were plotted using EnhancedVolcano (cutoff: logFCcutoff=0, pvalCutoff=0.05).

### Tissue culture

HEK-293T cells were cultured in Dulbecco’s Modified Eagle Medium (Gibco, #11960069) supplemented with 10% fetal bovine serum (FBS) (Life Technologies, #16141) and 1x Pen-Strep (#15140122, Life Technologies). SB28 mouse glioblastoma cells were cultured in RPMI with 10% FBS, and 1% HEPES, sodium pyruvate, NEAA, Pen-Strep, Glutamax, and beta-mercaptoethanol. GBM 43 cells were cultured in N5 media. Cell cultures were authenticated by short tandem repeat (STR) analysis at the UC Berkeley DNA Sequencing Facility and routinely tested for mycoplasma using the MycoAlert Detection Kit (Lonza, #75866-212) as part of standard practice.

For siRNA experiments, cells were transfected with jetOPTIMUS transfection reagent and buffer (Polyplus Transfection) using siRNA against *SHOC2*. Transfections were carried out according to jetOPTIMUS kit instructions for reverse transfection. Briefly, siRNAs were combined with jetOPTIMUS transfection buffer and reagent with a 1:1 ratio of nucleic acid:reagent and incubated at room temperature for 10 minutes. This master mix was then added to a 6-well plate prepared with normal N5 media. Cells were then split normally and added to the 6-well plate with 400,000 cells per well. Cells were then incubated as normal and siRNA knockdown of *SHOC2* was observed for 24-48 hours following transfection.

### Immunoblotting

Whole cell lysates were harvested using standard methods in RIPA buffer (50 mM Tris-HCl at pH 8.0, 150 mM NaCl, 0.5% Deoxycholate, 0.1% SDS, 1% IGEPAL CA630) with fresh protease (#P8340, Sigma) and phosphatase inhibitor (#P2850, Sigma) cocktails. A total of 15 μg of protein was loaded into pre-cast NuPAGE electrophoresis gels (Life Technologies). Samples were separated by SDS-PAGE, transferred to nitrocellulose or PVDF membranes, and blocked in either 5% bovine serum albumin or 5% skim milk in TBS buffer for 1 hour at room temperature. Primary antibodies were incubated overnight at the indicated dilutions at 4 degrees Celsius and HRP conjugated secondary antibodies were incubated for 1 hour at room temperature followed by ECL based detection on film. The following antibodies were used: phospho-ERK (Cell Signaling Technologies, #4370, 1:1000 dilution), total ERK (Cell Signaling Technologies, #4695, 1:1000 dilution), phospho-MEK (Cell Signaling Technologies, #9121, 1:1000 dilution), total MEK (Cell Signaling Technologies, #8727, 1:1000 dilution), and SHOC2 (Cell Signaling Technologies, #53600, 1:1000 dilution).

### CRISPRi cell line generation and genome-wide screening

Lentivirus containing pMH0001 (UCOE-SFFV-dCas9-BFP-KRAB, #85969, Addgene) was produced from transfected HEK293T cells with packaging vectors (pMD2.G #12259, Addgene, and pCMV-dR8.91, Trono Lab) following the manufacturers protocol (#MIR6605, Mirus). SB28 and GBM43 cells were stably transduced to generate parental dCas9-KRAB-BFP cells and selected by flow cytometry using a SH800 sorter (Sony). Subsequent gene specific knockdowns were achieved by individually cloning single-guide RNA (sgRNA) protospacer sequences into the pCRISPRia-v2 vector (#84832, Addgene) between BstXI and BlpI restriction sites. All constructs were validated by Sanger sequencing of the protospacer region. The following protospacers were used: sgNTC (GTGCACCCGGCTAGGACCGG), sghSHOC2-1 (GGGCAGCGTCGCTTCTTAGG), sghSHOC2-2 (GGGCTCCTGACGGTAACTCG).

For human GBM43 genome-wide CRISPRi screening, we used a compact and highly active sgRNA library containing the top 2 on-target sgRNAs for 23,483 genes that was optimized through aggregation of 126 genome-wide CRISPRi screens, established sgRNAs targeting essential genes, and machine learning prediction algorithms.^29^ This genome-wide dual sgRNA library has been previously validated through multiple growth-based screens as well as through confirmation of on-target gene repression using perturb-seq, exhibiting 82–92% median target knockdown. The genome-wide dual sgRNA library was cloned into the library expression vector pU6-sgRNA Ef1alpha Puro-T2A-GFP derived frompJR85 (#140095, Addgene) and modified to express a second sgRNA using the human U6 promoter as previously described.

1,137 non-targeting sgRNA pairs were also included as negative controls in the screen. To generate lentiviral pools, HEK293T cells were transfected with the sgRNA library along with packaging plasmids as described above, and viral supernatant was collected 72 h following transfection. Lentiviral libraries were infected into GBM43 dCas9-KRAB-BFP cells, cultured for 2 days following infection, selected in 0.5 μg/mL puromycin for 2 days, and then allowed to recover in standard growth media for 1 day. Infection efficiency was evaluated by measuring GFP positivity on flow cytometry, and cell pellets were subsequently frozen down at this “T0” timepoint. Cells were cultured in either 1 μM selumetinib or vehicle (DMSO) control for 16 days. Cell pellets were frozen down at this “T16” timepoint for both vehicle and selumetinib treated conditions. Samples were then processed for sgRNA abundance library preparation using Q5 High-Fidelity DNA Polymerase (NEB) and sequenced on an Illumina NextSeq-500.^30^ Downstream analysis was carried out as previously described.^20^ In brief, enrichment or depletion of sgRNA abundances were determined by down sampling trimmed sequencing reads to equivalent amounts across all samples, and then calculating the log2 ratio of sgRNA abundance in experimental conditions to sgRNA abundance in vehicle conditions at T16, or between sequencing reads from T16 and T0 timepoints within experimental or control conditions. Specifically, we computed normalized log2 ratios for selumetinib-treated sgRNA abundance at T16 compared to vehicle-treated sgRNA abundance at T16 to identify mediators of selumetinib responses. Statistical significance was calculated using Wald test comparing replicates across conditions without a log2 fold change threshold. Hits were prioritized by normalizing log2 ratios to the total number of population doublings in the screen and the standard deviations of the non-targeting control sgRNAs. These phenotype log2 ratios were used for subsequent analysis and visualization. Genes were filtered at an adjusted p-value < 0.05 for further analysis.

For mouse SB28 genome-wide CRISPRi screening, we used a genome-wide mouse sgRNA library targeting 20,003 genes at 5 sgRNAs/gene, for a total of 107,415 sgRNAs, in addition to 2170 non-targeting control sgRNAs.^31^ Pooled lentivirus was generated as above, and SB28 cells were transduced with the addition of polybrene (8 µg/mL). 1 day of puromycin selection (2.0 µg/mL) was performed, followed by 1 day of growth in non-puromycin 10% FBS in DMEM media. Two replicates of each screen were performed at a coverage of 1000x cells per sgRNA, in both vehicle and 1 μM selumetinib conditions. Infection efficiency was evaluated by measuring GFP positivity on flow cytometry. Initial (T0) cell populations were frozen in 10% DMSO and processed for genomic DNA using the NucleoSpin Blood XL Kit (Machery-Nagel, #740950.50). Endpoint cell pellets were harvested for genomic DNA after 10 days of growth, corresponding to ∼10 and ∼5 doublings in the vehicle and selumetinib conditions, respectively. sgRNA sequencing libraries were prepared using NEBNext Ultra II Q5 PCR MasterMix (New England Biolabs, #M0544L) and sequenced on an Illumina NextSeq-500.

Growth phenotype (gamma) was defined as log2(sgRNA count (vehicle) / sgRNA count T0) minus median sgNTC log2(sgRNA count (vehicle) / sgRNA count T0), then normalized by the number of cell doublings, as previously described.^20^ Drug phenotype (tau) was defined as log2(sgRNA count (selumetinib) / sgRNA count T0) minus median sgNTC log2(sgRNA count (selumetinib) / sgRNA count T0), then normalized by the number of cell doublings in the drug screen. Drug:growth ratio phenotype (rho) was defined as log2(sgRNA count (selumetinib) / sgRNA count (vehicle)), then normalized by the number of cell doublings in the drug screen.

Gene-level phenotypes were summarized as the mean of the top 3 sgRNAs against a given gene, ranked according to screen phenotype. Statistical significance was calculated using Mann-Whitney-U test for a given perturbation compared to the sgRNA distribution of the non-targeting control sgRNAs.

### Nucleic acid extraction and qRT-PCR

From glioblastoma cell lines, RNA was extracted using the RNeasy Mini Kit (#74106, QIAGEN) according to manufacturer’s instructions. cDNA was synthesized from RNA using iScript cDNA Synthesis kit (#1708891, BioRad). Real-time QPCR was performed using PowerUp SYBR Green Master Mix (#A25918, Thermo Fisher Scientific) on a QuantStudio 6 Flex Real Time PCR system (Life Technologies) and analyzed using the double delta method as previously reported.^20^ The following QPCR primers were used: GAPDH-F (5′- GTCTCCTCTGACTTCAACAGCG-3′), GAPDH-R (5′-ACCACCCTGTTGCTGTAGCCAA-3′), hSHOC2-F (5’- GTTGACAATACGATCAAACGGC-3’), hSHOC2-R (5’- CTCTTCCCGGCATTTGTTGAG-3’), mSHOC2-F (5’-AATACCATCAAACGGCCAAATCC-3’), and mSHOC2-R (5’- AACCGCATTGAGTTCTCCTCC-3’).

## Data Availability Statement

Human tumor DNA methylation (n=129), single nuclear RNA sequencing (n=9) and intracranial tumor single cell RNA-sequencing (n=6) reported in this manuscript have been deposited in the NCBI Gene Expression Omnibus under Bioproject PRJNA111929.

## Code availability

The open-source software, tools, and packages used for data analysis in this study, as well as the version of each program, were ImageJ (v2.1.0), R (v3.5.3 and v3.6.1), cellranger (v6.1.2), Seurat R package (v4.4.0), Clustree (v0.5.0), Scanorama (v1.7.3), minfi (Bioconductor v3.10), ConsensusClusterPlus (Bioconductor v3.10), Heatmap.2 R package (gplots v3.13), and ggplot2 (v3.4.3). No custom software, tools, or packages were used. CRISPRi screen analysis code is available at https://github.com/liujohn/CRISPRi-dual-sgRNA-screens/blob/main/module2/PhenotypeScores.R.

---

**Supplementary Figure 1.**
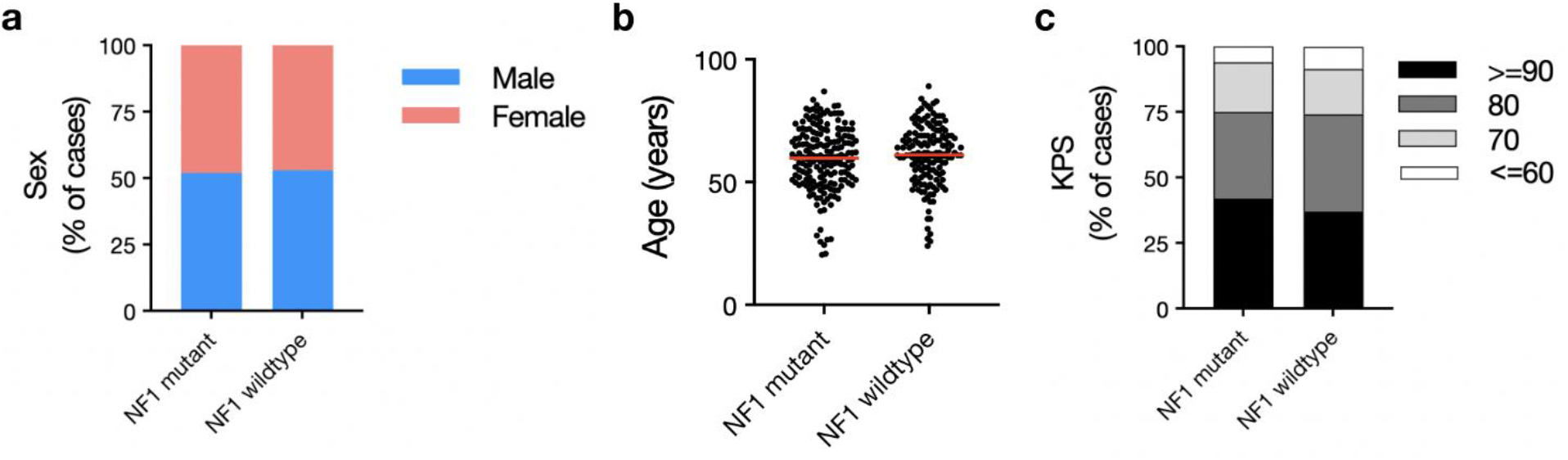
Paired analysis of the *NF1* mutant, IDH-wildtype glioblastoma cohort with a propensity score matched *NF1* wildtype, IDH-wildtype glioblastoma cohort. Comparison of baseline clinical parameters between *NF1* mutant and *NF1* wildtype propensity score matched cohort shows no differences in a. sex, b. age, or c. KPS.

**Supplementary Figure 2.**
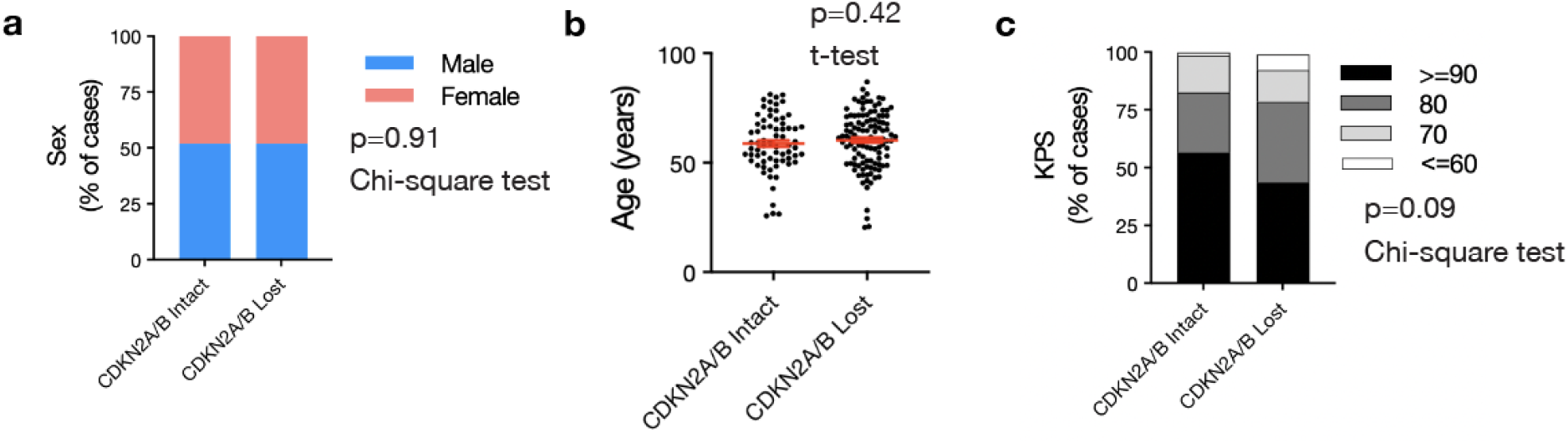
*CDKN2A/B* status in *NF1* mutant, IDH-wildtype glioblastomas and relationship to baseline clinical or pathologic variables. a. No significant difference was observed in biological sex (p=0.91,Chi-square test), b. age (p=0.42, unpaired t-test) or c. KPS (p=0.09, Chi-Square test) based on *CDKN2A/B* status.

**Supplementary Figure 3.**
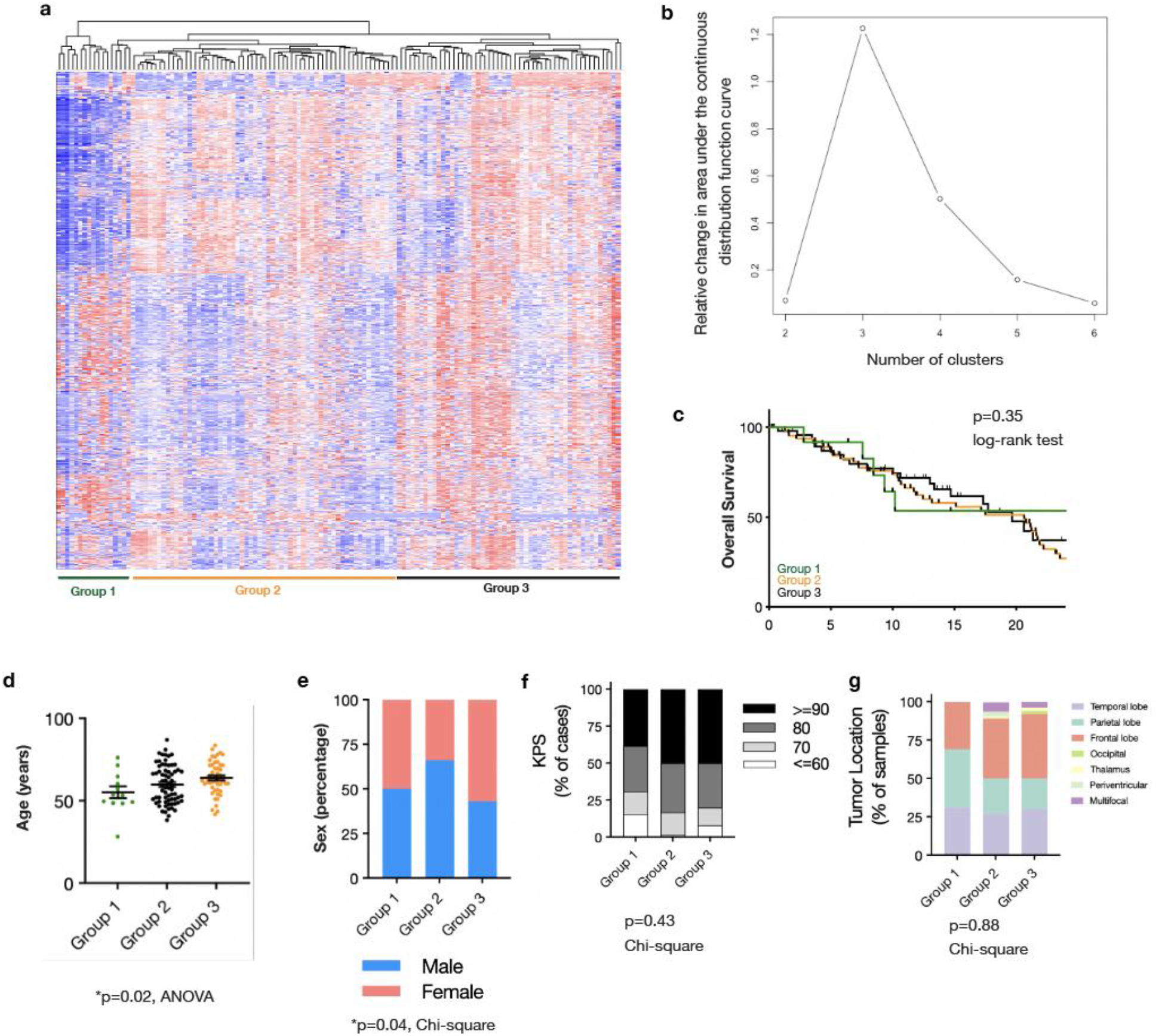
DNA methylation array analysis of *NF1* mutant, IDH-wildtype glioblastomas identifies 3 epigenetic groups. a. Hierarchical clustering identifies 3 epigenetic *NF1* mutant glioblastoma groups. b. Change in area under the curve for consensus cumulative distribution function (CDF) reveals greater than 3 clusters results in minimal appreciable increase. c. DNA methylation groups stratified by clinical variables showed no statistically significant difference in OS. d. DNA methylation groups show significant differences in age (p=0.02, ANOVA) and e. biologic sex (p=0.04, Chi-square) but are not significantly different with regard to f. KPS (p=0.43, Chi-square test), or g. tumor location (p=0.88, Chi-square test).

**Supplementary Figure 4.**
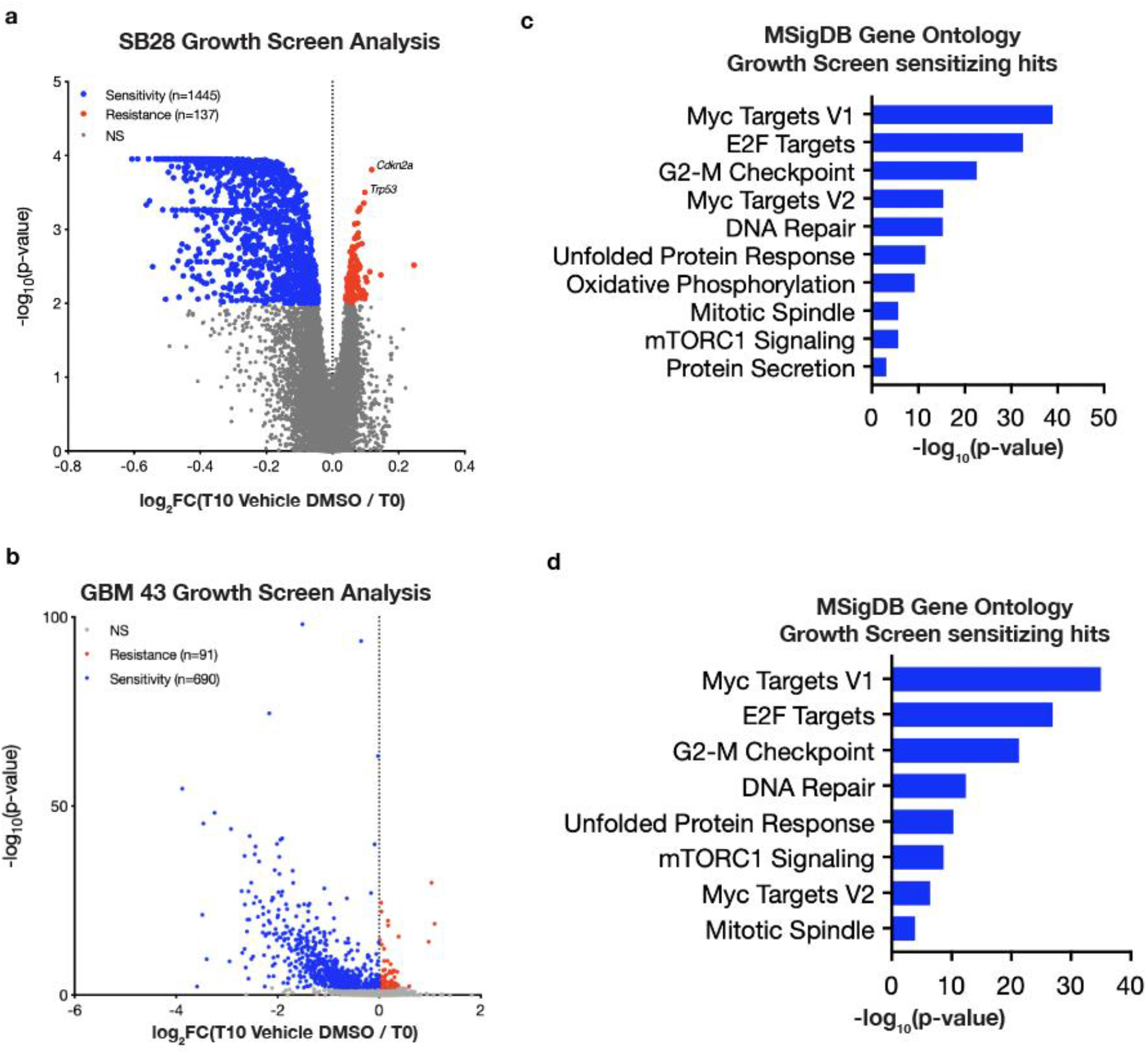
Genome-wide CRISPRi screens for mediators of cell growth in SB28 or GBM43 glioblastoma cells reveals a conserved requirement for cell cycle genes. a. Volcano plot of genes significantly mediating cell growth (T10 Vehicle DMSO / T0) reveals 1445 genes underlying decreased growth and 137 genes underlying increased growth. Notably, *Cdkn2a* or *Tp53* repression are the top hits leading to increased cell growth. b. Volcano plot of genes significantly mediating cell growth (T10 Vehicle DMSO / T0) reveals 690 genes underlying decreased growth and 91 genes underlying increased growth. c. Gene ontology analysis of genes leading to decreased glioblastoma cell growth when repressed reveals enrichment cell cycle gene sets (E2F Targets, G2-M Checkpoint, DNA Repair, Mitotic Spindle). d. Gene ontology analysis of genes leading to decreased glioblastoma cell growth when repressed reveals enrichment cell cycle gene sets (E2F Targets, G2-M Checkpoint, DNA Repair, Mitotic Spindle) consistent with SB28 CRISPRi growth screen hits.

**Supplementary Figure 5.**
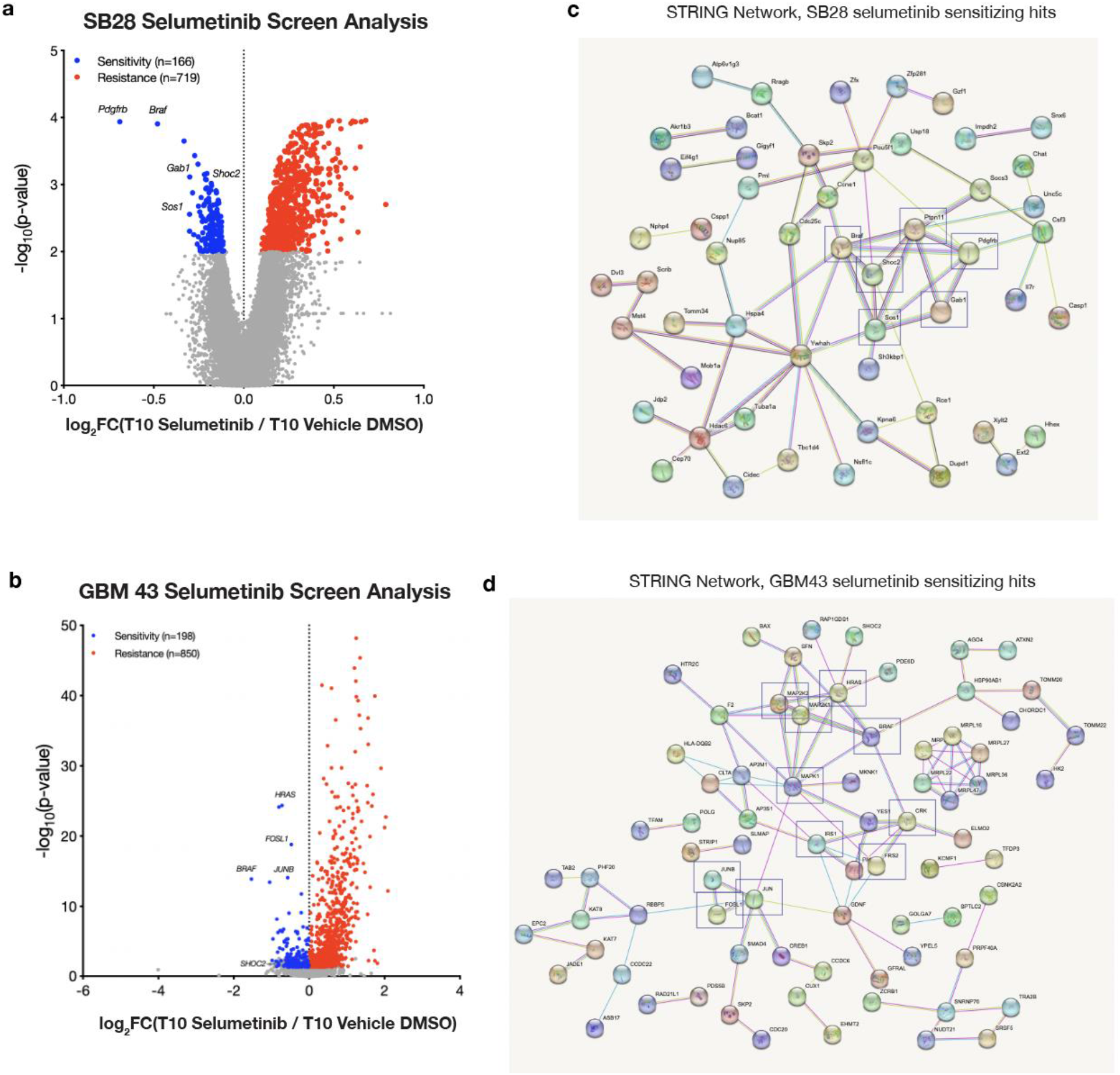
Genome-wide CRISPRi screen in GBM43 human glioblastoma cells reveals cell cycle genes are required for growth and Ras/Raf/MEK/ERK signaling components underlie MEK inhibitor sensitivity. a. Volcano plot of genes significantly mediating response to the MEK inhibitor selumetinib (T10 Selumetinib / T10 Vehicle DMSO control) reveals 166 genes mediating sensitivity and 719 genes mediating resistance. b. Protein- protein interaction network (STRING) shows genes mediating sensitivity are enriched for Ras/Raf/MEK/ERK pathway genes such as *Braf, Pdgfrb, Ptpn11, Shoc2, Sos1,* and *Gab1.* c. Volcano plot of genes significantly mediating response to the MEK inhibitor selumetinib (T10 Selumetinib / T10 Vehicle DMSO control) reveals 198 genes mediating sensitivity and 850 genes mediating resistance. d. Consistent with SB28 CRISPRi selumetinib screen hits, protein- protein interaction network (STRING) shows genes mediating sensitivity are enriched for Ras/Raf/MEK/ERK pathway genes such as *HRAS, BRAF, MAP2K1, MAP2K2, SHOC2, MAPK1, CRK, FRS2, IRS1, JUN, JUNB,* and *FOSL1*.

**Supplementary Figure 6.**
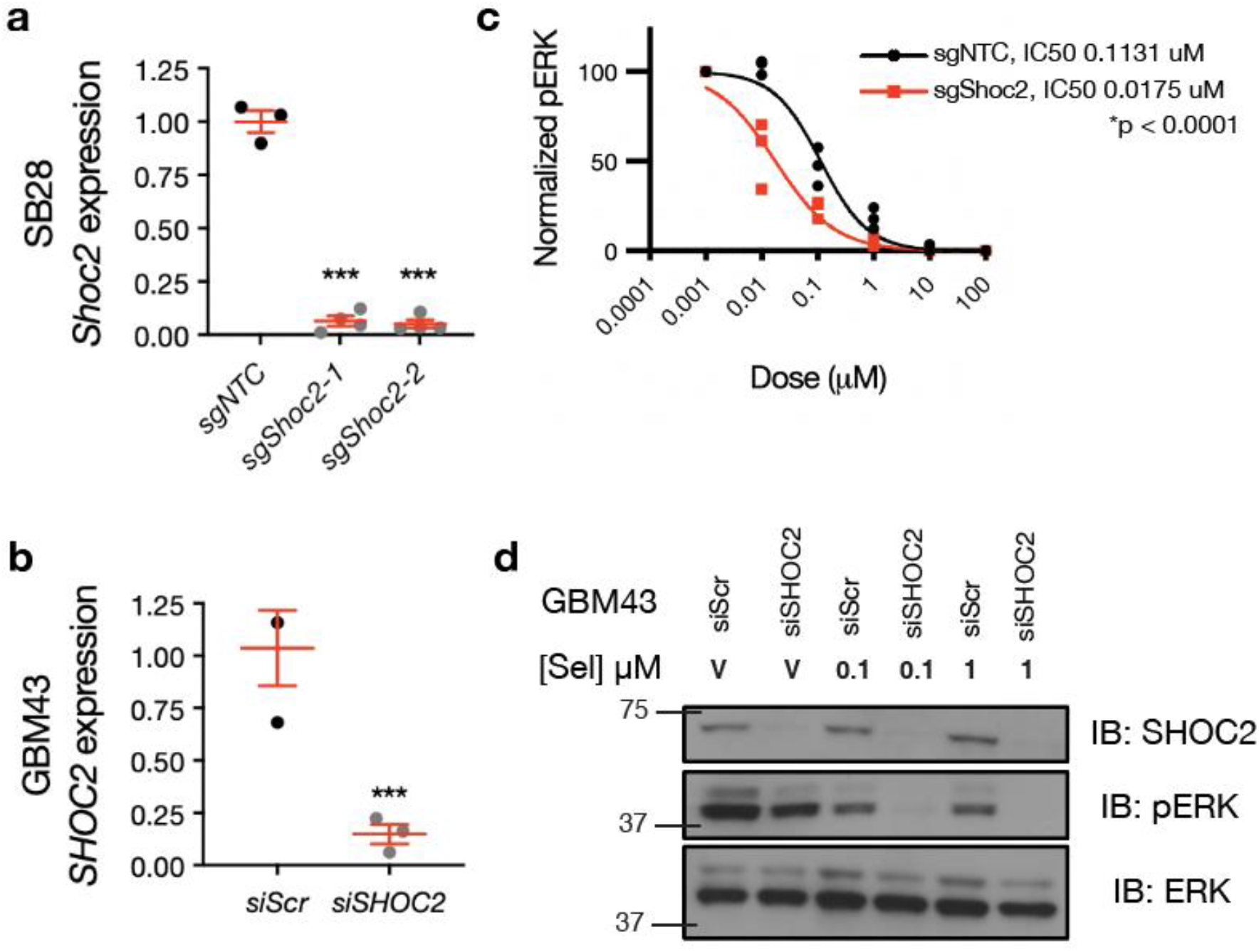
Validation and analysis of *SHOC2* deficient glioblastoma cells. a. QRT-PCR Validation of CRISPRi sg*Shoc2* SB28 cells. b. QRT-PCR validation of siRNA *siSHOC2* GBM43 cells. c. Quantification of selumetinib response in sg*Shoc2* deficient SB28 glioblastoma cells (IC50 0.0751, 95% CI 0.011 – 0.027) versus sgNTC SB28 glioblastoma cells (IC50 0.1131, 95% CI 0.080 – 0.169) (p<0.0001, F-test). d. Immunoblot validation of si*SHOC2* GMB43 cells and selumetinib dose response shows increased sensitivity in si*SHOC2* deficient cells.

**Supplementary Figure 7.**
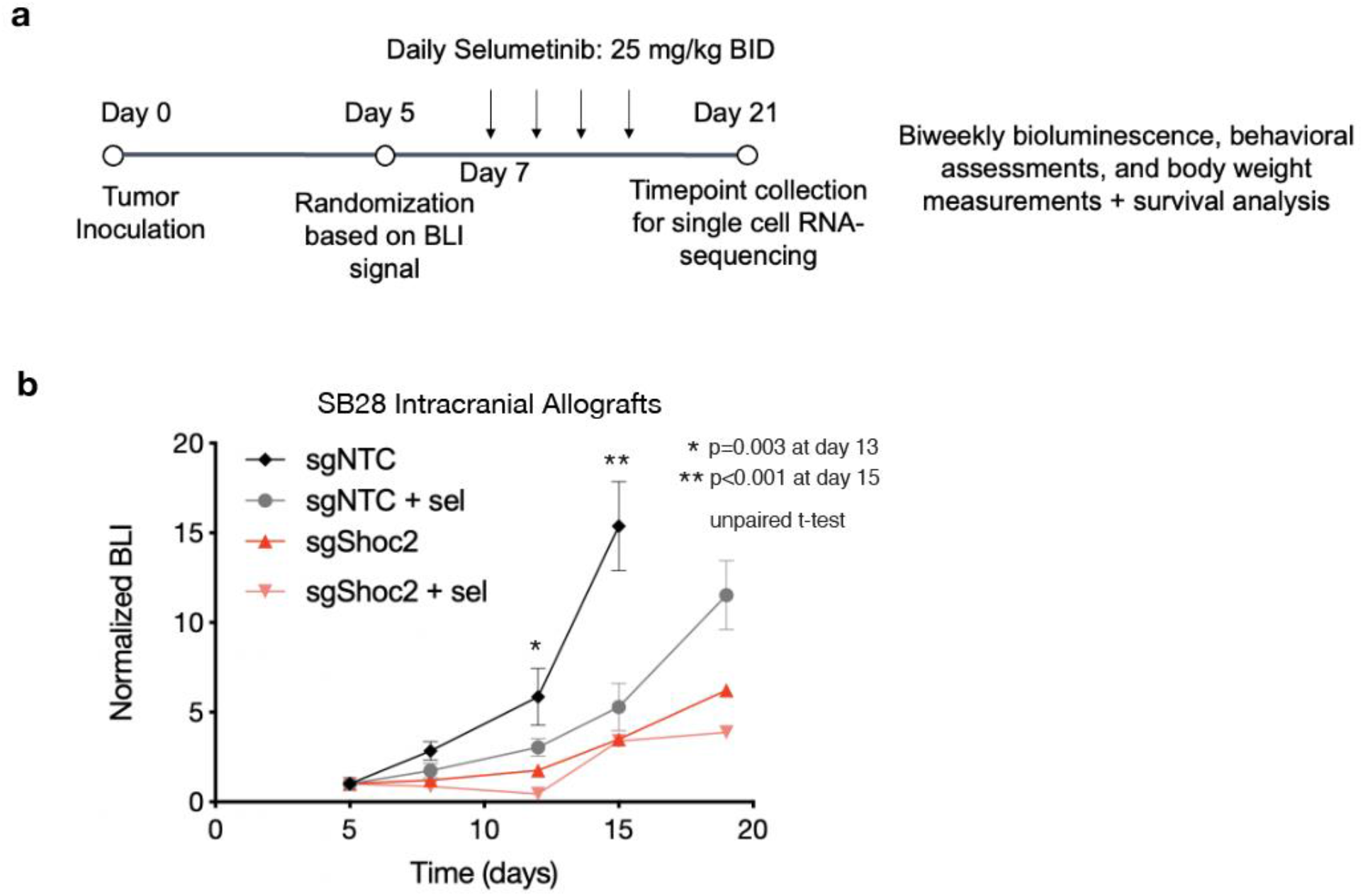
Normalized bioluminescence (BLI) analysis following oral selumetinib treatment in CRISPRi control sgNTC or CRISPRi sg*Shoc2* SB28 allografts. a. Overview of mouse SB28 intracranial allograft experiment in immunocompetent host mice. b. BLI reveals *Shoc2* loss impedes tumor growth and is sufficient to further sensitize *NRAS^G12V^* mutant SB28 intracranial allografts to selumetinib (unpaired t-test).

**Supplementary Figure 8.**
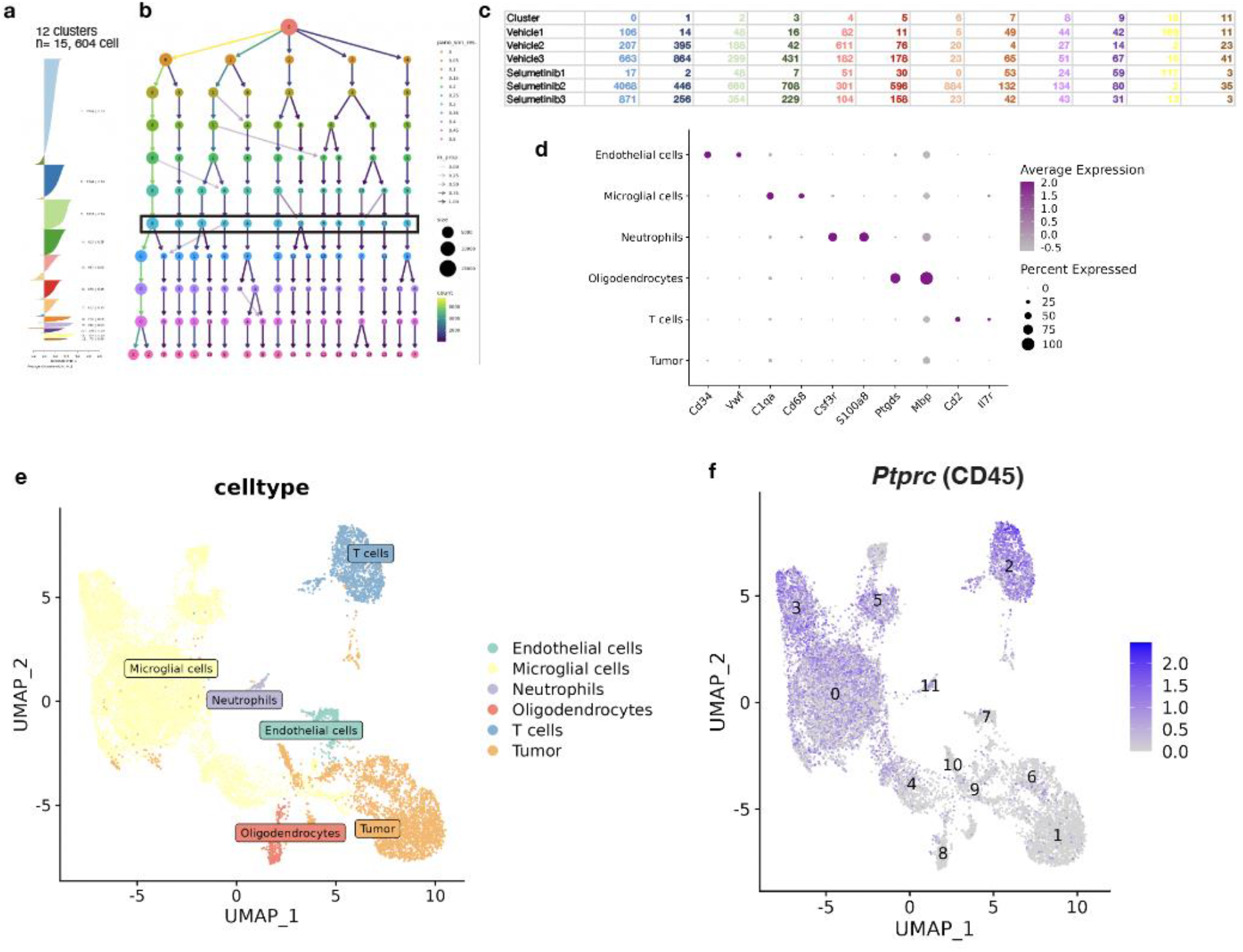
Single cell RNA-sequencing of SB28 intracranial allografts reveals 12 clusters comprised of both tumor and non-tumor cells. a. Silhouette analysis and b. cluster tree reveals 12 clusters leads to robust cell type classification. c. Summary of number of cells per cluster by sample of origin. d. Marker gene expression and e. UMAP with cell type assignment for all detected cells separates tumor and non-tumor cell populations. f. *Ptprc* (CD45) expression marks non-tumor cells from the hematopoietic lineage. Non-tumor cells were filtered from further analysis.

**Supplementary Figure 9.**
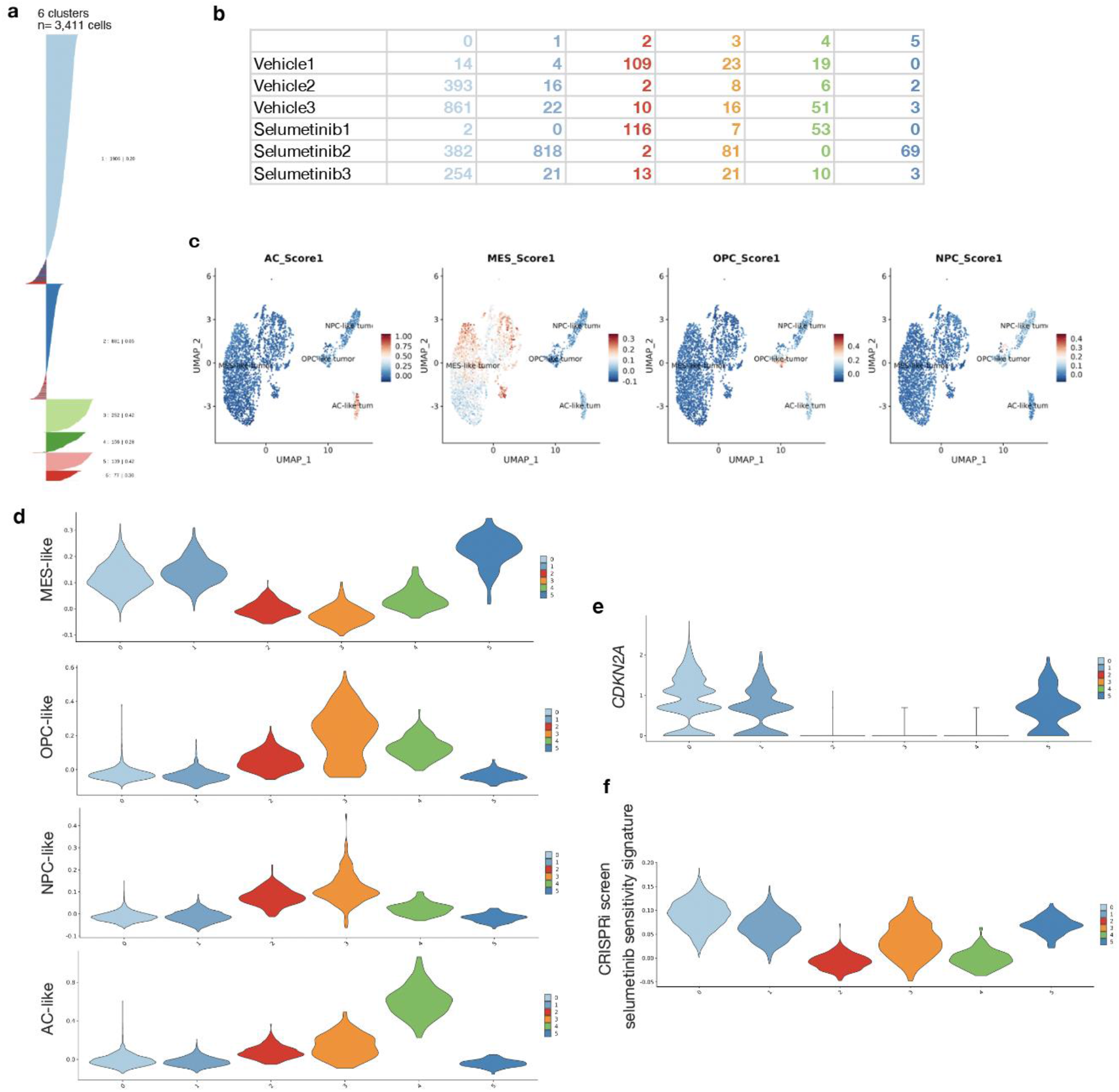
Single cell RNA-sequencing analysis of tumor cells from SB28 allografts treated with selumetinib reveals the MES-like subpopulation is associated with *CDKN2A* retention and selumetinib sensitivity. a. Silhouette analysis reveals 6 clusters leads to robust cell type classification. b Summary of number of cells per tumor cell cluster by sample of origin. c. Feature plots for MES-like, OPC-like, AC-like, and NPC-like cell state signature expression scores. d. Violin plots for MES-like, OPC-like, AC-like, and NPC-like cell state signature expression scores for cluster type assessment. e. *CDKN2A* and f. CRISPRi screen selumetinib sensitivity signature feature violin plots reveal correlation with MES-like cell state.

**Supplementary Figure 10.**
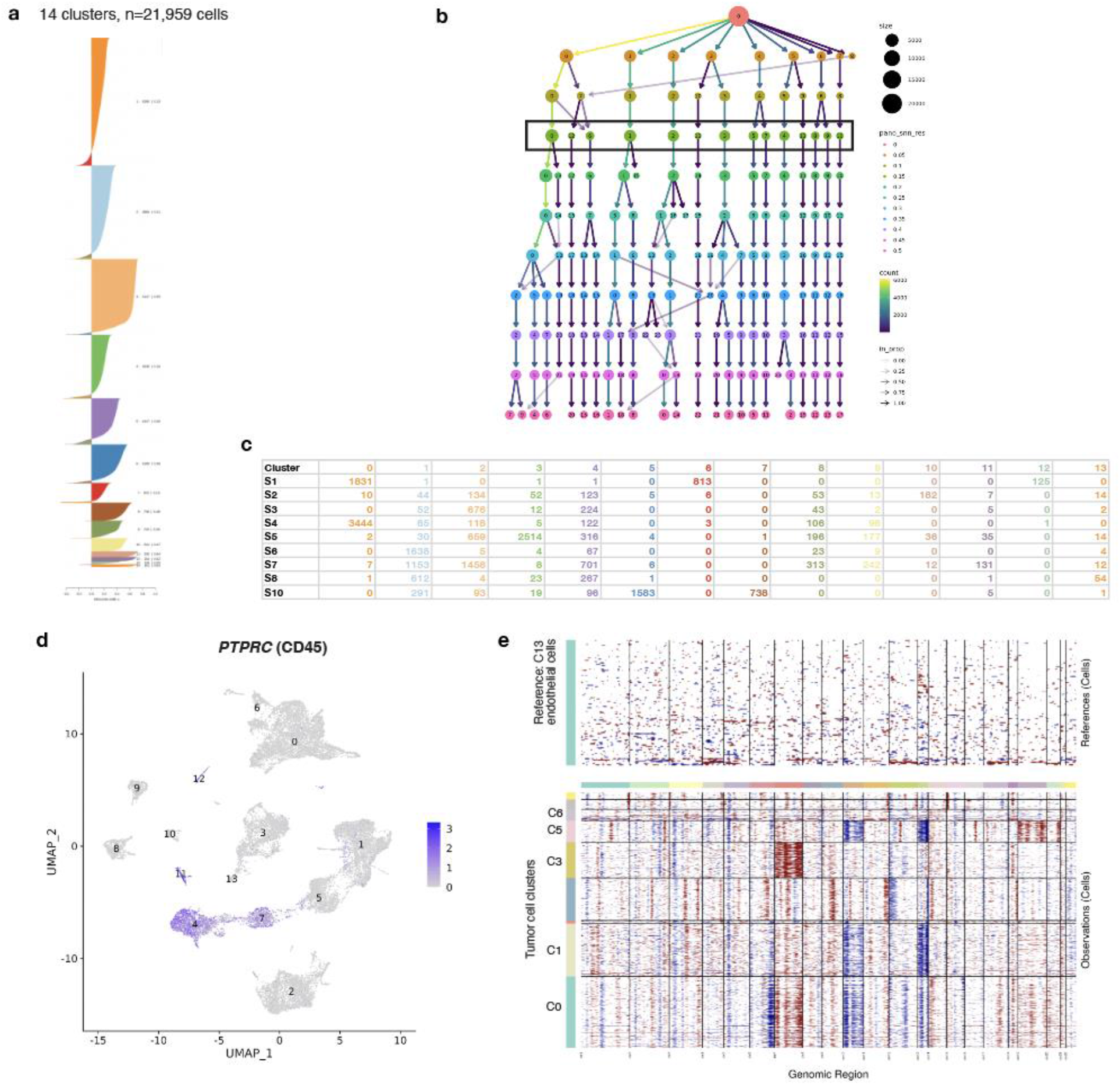
Single nuclear RNA-sequencing analysis of human *NF1* mutant, IDH-wildtype glioblastomas (n=9). a. Silhouette analysis and b. cluster tree reveals 14 clusters across 21,959 single nuclei leads to robust cell type classification. c. Summary of number of cells per cluster by sample of origin. d. *PTPRC* (CD45) expression marks non-tumor cells from the hematopoietic lineage. e. inferCNV identifies tumor cell clusters (clusters 0, 1, 3, 5, 6,) containing CNAs in single cell RNA-seq data compared to non-tumor populations with normal relative ploidy.

**Supplementary Figure 11.**
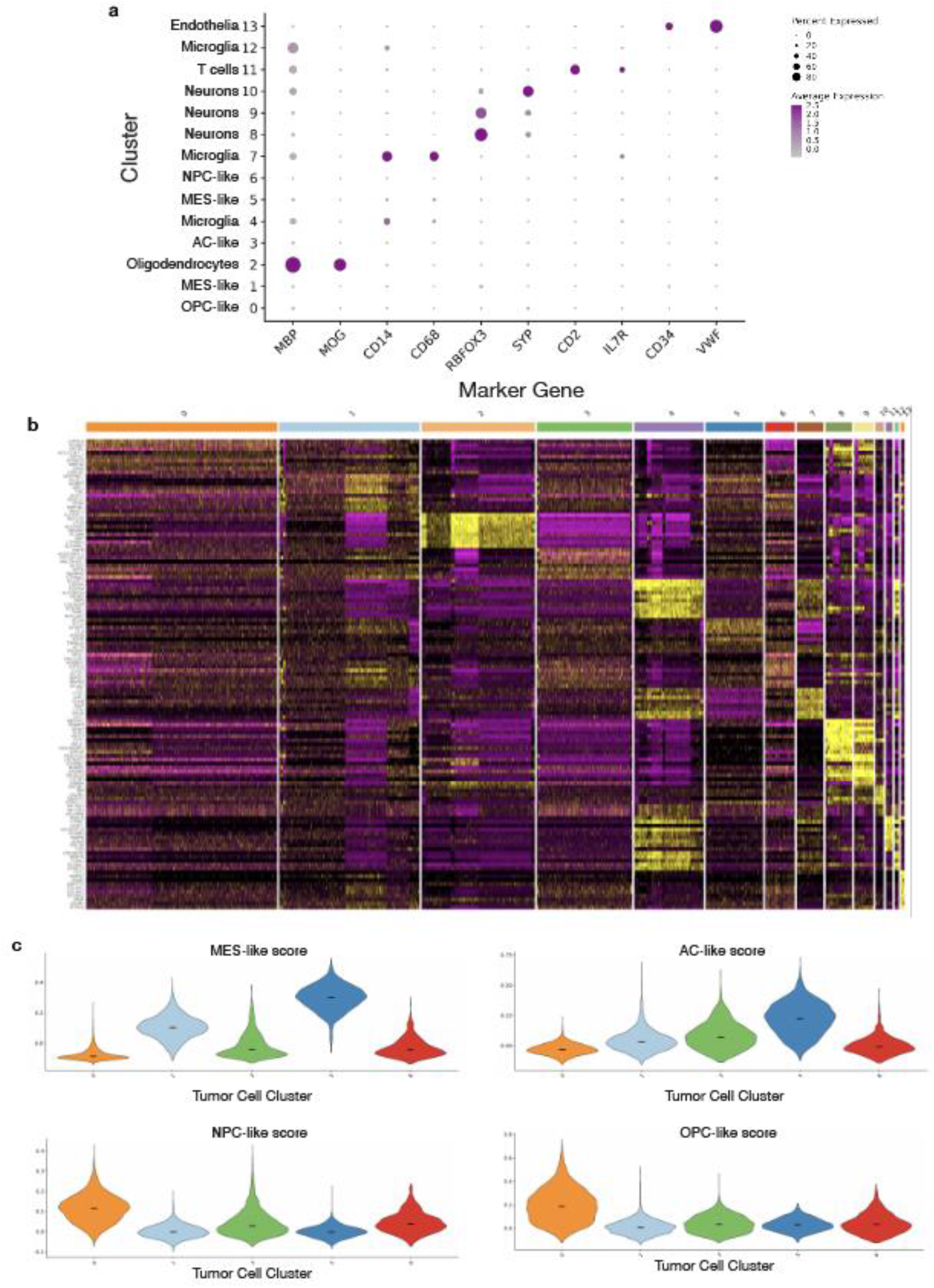
Marker gene analysis and cluster assignment from snRNA-sequencing of human *NF1* mutant, *CDKN2A/B* deleted glioblastomas. a. Dot plot for non-tumor cell population marker gene expression. b Heatmap for cluster marker genes across all clusters. c. Violin plots for MES-like, OPC-like, AC-like, and NPC-like cell state signature expression scores in tumor cell clusters.

## Supplementary Tables

**Supplementary Table 1.** Compiled variant list from the UCSF500 assay, a CLIA certified, capture based targeted DNA sequencing assay of 529 cancer-associated genes across *NF1* mutant, IDH-wildtype glioblastomas (n=186).

**Supplementary Table 2.** Propensity score matching to identify a *NF1* wildtype, IDH-wildtype glioblastoma cohort for baseline comparison.

**Supplementary Table 3.** Univariable Cox proportional hazards analysis using recurrently co-mutated genes across NF1 mutant and NF1 wildtype glioblastoma cohorts.

**Supplementary Table 4.** Univariable and multivariable Cox proportional hazards analysis using clinical factors and CDKN2A/B mutation across NF1 mutant glioblastomas.

**Supplementary Table 5.** Normalized beta values for top 5000 most variable DNA methylation probes across *NF1* mutant, IDH wildtype glioblastomas (n=129).

**Supplementary Table 6.** Analysis of GBM43 CRISPRi screen comparing sgRNA enrichment between cells treated with selumetinib for 10 days compared to treated with vehicle for 10 days.

**Supplementary Table 7.** Analysis of SB28 CRISPRi screen comparing sgRNA enrichment between cells treated with selumetinib for 10 days compared to treated with vehicle for 10 days.

**Supplementary Table 8.** Cluster marker gene list for scRNA-sequencing of tumor cells from selumetinib (n=3) or vehicle treated (n=3) SB28 intracranial glioblastomas.

**Supplementary Table 9.** Differential gene expression analysis between selumetinib and vehicle treated tumor cells from SB28 intracranial glioblastomas.

**Supplementary Table 10.** Cluster marker gene list for snRNA-sequencing of human NF1 mutant glioblastomas (n=9).

